# ConDoR: Tumor phylogeny inference with a copy-number constrained mutation loss model

**DOI:** 10.1101/2023.01.05.522408

**Authors:** Palash Sashittal, Haochen Zhang, Christine A. Iacobuzio-Donahue, Benjamin J. Raphael

## Abstract

Tumors consist of subpopulations of cells that harbor distinct collections of somatic mutations. These mutations range in scale from single nucleotide variants (SNVs) to large-scale copy-number aberrations (CNAs). While many approaches infer tumor phylogenies using SNVs as phylogenetic markers, CNAs that overlap SNVs may lead to erroneous phylogenetic inference. Specifically, an SNV may be lost in a cell due to a deletion of the genomic segment containing the SNV. Unfortunately, no current single-cell DNA sequencing (scDNA-seq) technology produces accurate measurements of both SNVs and CNAs. For instance, recent *targeted* scDNA-seq technologies, such as Mission Bio Tapestri, measure SNVs with high fidelity in individual cells, but yield much less reliable measurements of CNAs. We introduce a new evolutionary model, the *constrained k-Dollo model*, that uses SNVs as phylogenetic markers and partial information about CNAs in the form of clustering of cells with similar copy-number profiles. This copy-number clustering constrains where loss of SNVs can occur in the phylogeny. We develop ConDoR (Constrained Dollo Reconstruction), an algorithm to infer tumor phylogenies from targeted scDNA-seq data using the constrained *k*-Dollo model. We show that ConDoR outperforms existing methods on simulated data. We use ConDoR to analyze a new multi-region targeted scDNA-seq dataset of 2153 cells from a pancreatic ductal adenocarcinoma (PDAC) tumor and produce a more plausible phylogeny compared to existing methods that conforms to histological results for the tumor from a previous study. We also analyze a metastatic colorectal cancer dataset, deriving a more parsimonious phylogeny than previously published analyses and with a simpler monoclonal origin of metastasis compared to the original study.

**Code availability:** Software is available at https://github.com/raphael-group/constrained-Dollo

## Background

Cancer is an evolutionary process in which somatic mutations across all genomic scales – ranging from single nucleotide variants (SNVs) to large-scale copy number aberrations (CNAs) – accumulate in a population of cells. This process results in a heterogeneous tumor with subpopulations of cells, called *clones*, with distinct genomes. Reconstruction of the evolutionary history of cancer clones, known as a tumor phylogeny, from genomic sequencing data of the cells in a tumor is crucial for understanding cancer progression and developing effective therapies for treatment [1–4].

Early cancer sequencing projects performed bulk sequencing of tumor samples and thus measured somatic mutations from a mixture of thousands or millions of cells. Tumor phylogeny inference from this data is complicated since it requires deconvolution of the data, i.e. simultaneous inference of the tumor clones and their proportions in the mixture [5–11]. Recent developments in single-cell DNA sequencing (scDNA-seq) allow parallel sequencing of thousands of individual cells from a tumor [2, 12–15], alleviating the need for such deconvolution. However, tumor phylogeny inference from this data remains challenging since current scDNA-seq technologies are error-prone and produce data with missing information. As such, phylogeny inference using scDNA-seq data involves correcting these errors and imputing the missing data under some evolutionary model [2, 16].

Multiple evolutionary models have been used to construct tumor phylogenies from scDNA-seq data. Early works [17–21] used SNVs as evolutionary markers, and relied on the *infinite-sites model* [22] which states that an SNV can be gained only once and never be subsequently lost in the phylogeny. While the same SNV occurring independently more than once is rare [23], loss of SNVs due to copy-number deletions is common in cancer [24]. To account for these losses, other works [25–28] use some variant of the *k-Dollo model* [29], in which a mutation can be gained at most once but may be lost at most *k* times during the course of the evolution, where *k* is a user-defined integer. Several methods [30, 31] employ an even more permissive model, the *finite-sites model* [32], which allowed mutations to be gained and lost multiple times.

A major limitation of the aforementioned models and methods is that they do not utilize any information about CNAs, which can often also be derived from scDNA-seq data. This limitation is addressed in methods such as SCARLET [33], BiTSC^2^ [34] and COMPASS [35] which incorporate copy-number information during phylogeny inference. SCARLET introduced a novel loss-supported Dollo model that requires the copy-number profile of each cell and the copynumber phylogeny as input. BiTSC^2^ and COMPASS, on the other hand, construct a joint phylogeny with both SNV and CNA events. However, these methods rely heavily on accurate and simultaneous identification of SNVs and CNAs on the same set of cells, which is challenging with the current scDNA-seq technologies [36].

Current scDNA-seq technologies fall into one of two classes with different capabilities from measuring CNAs and SNVs. First, *whole genome* scDNA-seq technologies yield data with roughly uniform coverage of the whole genome but with low depth at any particular locus, making it suitable for detection of larger CNAs in single-cells but not SNVs [12, 14, 37]. In contrast, *targeted* scDNA-seq technologies sequence specific regions of the genome, typically comprising of cancer-related genes, with high depth allowing accurate identification of SNVs but not of CNAs [13, 15, 25, 38]. For example, the Mission Bio Tapestri platform [39, 40] performs high-coverage sequencing (∼ 50× coverage) of hundreds of amplicons from thousands of cells. While precise identification of CNAs in each cell using such targeted scDNA-seq data is challenging, clustering of cells based on their copy-number profiles is a much simpler task. However, no existing evolutionary model utilizes such clustering information.

Here, we introduce a new evolutionary model, the *constrained k-Dollo model*, which allows losses of SNVs but constrains these losses to conform to a given copy-number clustering of cells (Figure 1). The key idea underpinning the constrained *k*-Dollo model is that, since loss of SNVs predominantly occurs due to CNAs, we allow loss of an SNV only between cells that have distinct copy-number profiles. Importantly, the constrained *k*-Dollo model generalizes both the infinite-sites and the *k*-Dollo models. Additionally, while loss of single nucleotide polymorphisms (SNPs), i.e. germline variants present in normal cells, can be informative during phylogeny inference, most existing methods only focus on somatic variants (SNVs) [17, 20, 26, 33]. We introduce the *Constrained k-Dollo Phylogeny Problem for Read Counts* (C*k*DP-RC) to infer a tumor phylogeny using scDNA-seq data, comprising of both SNVs and SNPs, and a clustering of the cells based on their copy-number profiles as input. We prove that the C*k*DP-RC problem is NP-hard and introduce ConDoR (Constrained Dollo Reconstruction), an algorithm that solves it exactly using mixed-integer linear programming.

**Figure 1:**
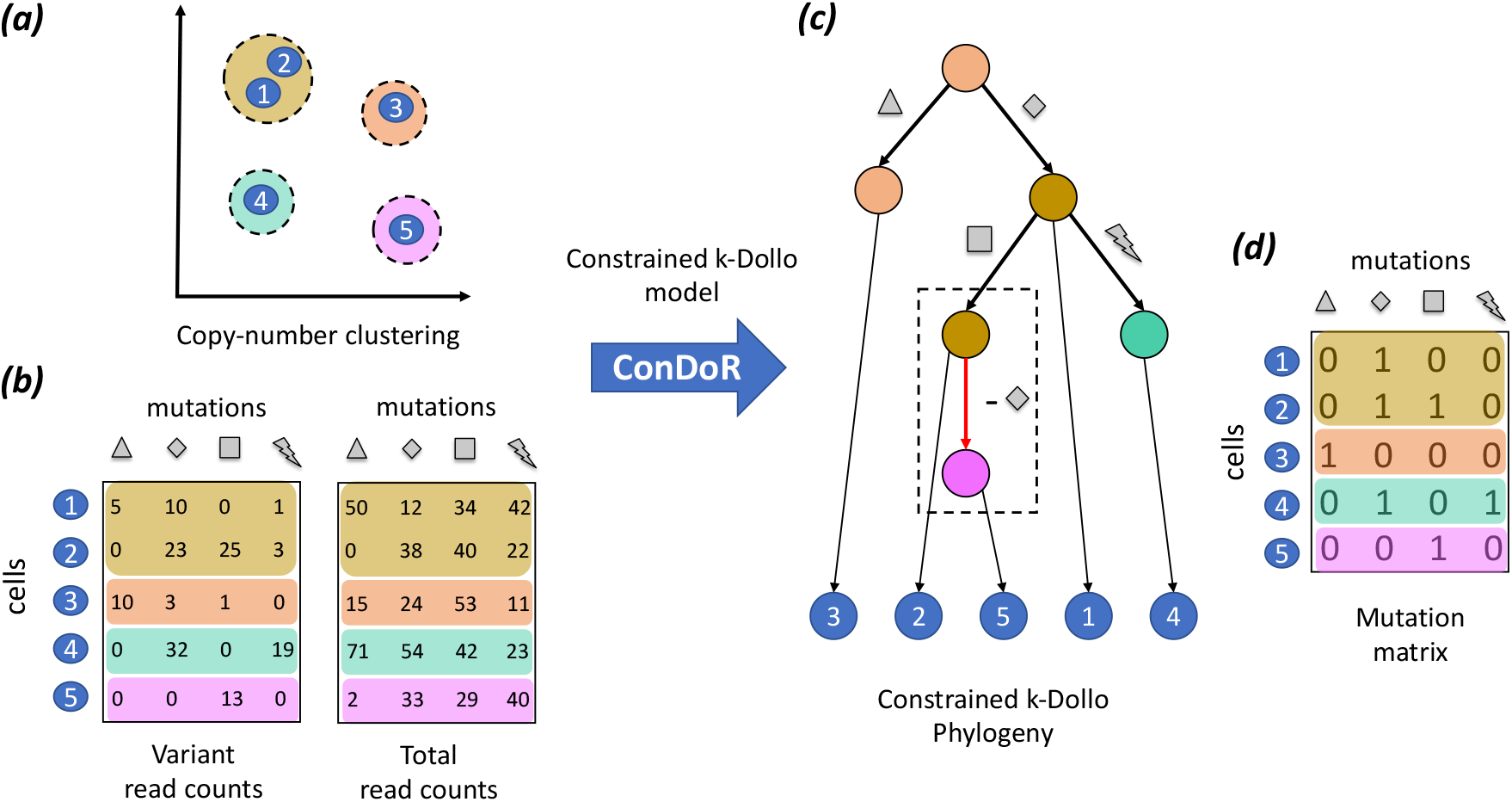
Overview of the ConDoR algorithm. ConDoR takes as input: (a) A clustering of cells based on copy-number profiles and (b) variant and total read counts from scDNA-seq data. ConDoR employs the Constrained k-Dollo model to construct the (c) constrained k-Dollo phylogeny with mutation losses (dashed box) allowed only between cells from distinct copy-number clusters and the (d) mutation matrix.

We show that ConDoR outperforms existing tumor phylogeny inference methods on simulated and real targeted scDNA-seq data. Specifically, ConDoR analysis of targeted scDNA-seq data from multiple regions of a pancreatic tumor results in a more plausible phylogeny compared to existing methods. Moreover, the ConDoR phylogeny provides insights into the evolution and spatial clonal architecture of the tumor which are supported by previous histopathological analysis of the tumor [41]. Second, ConDoR analysis of a metastatic colorectal cancer dataset from [38] demonstrates that the liver metastasis was seeded by the migration of a single cancer clone from the primary tumor to the metastasis (monoclonal seeding), in contrast to the more complicated polyclonal seeding reported in the original publication. At the same time, ConDoR obtains a more plausible explanation for the loss of mutations than other recent analyses of this data [20, 33].

## Results

### Constrained *k*-Dollo model

We propose a new model, the *constrained k-Dollo model*, that integrates information about SNVs, SNPs and CNAs on the same set of cells during phylogeny inference. Our model incorporates CNAs via a clustering of cells, where all cells in the same cluster have the same copy-number profile (set of CNAs). In other words, each cluster corresponds to a *copy-number clone*. We will refer to this clustering as the *copy-number clustering*.

Suppose we measure *m* SNVs and SNPs in *n* cells from a tumor. In the following, we collectively refer to SNVs and SNPs as mutations. We encode the presence or absence of mutations in the cells by an *n* × *m* binary *mutation matrix A* where *a*_*i,j*_ = 1 if cell *i* contains mutation *j* and *a*_*i,j*_ = 0 indicates the mutation *j* is absent in cell *i*. A phylogenetic tree *T* for the tumor is a rooted node-labeled tree which describes the evolutionary history of the tumor. Each internal node *v* in the tree *T* represents an ancestral cell and is labeled by a vector *a*_*v*_ ∈ {0, 1} ^*m*^ indicating the presence/absence of each mutation *j* ∈ [*m*], where [*m*] denotes the set {1, …, *m*}, in that cell. The root represents the normal cell and as such *a*_*r*(*T*),*j*_ = 0 if mutation *j* is an SNV and *a*_*r*(*T*),*j*_ = 1 if mutation *j* is an SNP. Each leaf of *T* corresponds to one of the *n* cells in the tumor. Our goal is to reconstruct a phylogenetic tree *T* for a given mutation matrix *A* under a given evolutionary model.

An edge (*v, w*) of a phylogeny *T* induces the *gain* of a mutation *j* ∈ [*m*] if *a*_*v,j*_ = 0 and *a*_*w,j*_ = 1. On the other hand, a mutation *j* ∈ [*m*] is said to be *lost* on edge (*v, w*) if *a*_*v,j*_ = 1 and *a*_*w,j*_ = 0. The simplest evolutionary model used in cancer genomics is the *infinite-sites model* which has two constraints [22]. Firstly, a mutation is allowed to be gained at most once in the phylogeny. This constraint stems from the *infinite-sites assumption* which posits that it is very unlikely for the same position in the genome to get mutated multiple time independently. Secondly, once a mutation is gained it cannot be subsequently lost. A phylogeny that satisfies these constraints is known as a perfect phylogeny [42].

While violation of the infinite-sites assumption is rare in cancer, SNVs and SNPs are frequently lost to due copynumber aberrations. As such, more recent phylogeny inference methods [26, 33] apply some variant of the *Dollo model* [29] for phylogeny inference, which allows loss of SNVs/SNPs. Specifically, under the Dollo model a mutation is allowed to be gained at most once but be lost multiple times in the phylogeny. The parameterized version of this model is the *k*-Dollo model, in which a mutation can only be lost at most *k* times in the phylogeny. However, a major limitation of Dollo models is that, although they allow loss of SNVs and SNPs, possibly due to CNAs, they do not incorporate any information about the copy-number states of the cells.

We introduce the constrained *k*-Dollo model that supplements the *k*-Dollo model with two additional constraints using the copy-number clustering of the cells. First, we assume that SNVs and SNPs can only be lost due to overlapping CNAs. As such, we only allow such losses between cells that belong to distinct copy-number clusters. Second, we assume that each copy-number clone arises only once in the phylogeny. As such, cells belonging to the same cluster form a connected subtree in the phylogeny. Let *p* be the number of copy-number clones and *σ* : [*n*] → [*p*] be the copy-number clustering of the *n* cells. We formally define the *constrained k-Dollo phylogeny* for a mutation matrix *A* and copy-number clustering *σ* as follows.

#### Definition 1

(constrained *k*-Dollo phylogeny). A *constrained k-Dollo phylogeny T* has the following properties.

1. Each node *v* ∈ *V* (*T*) is labeled by *a*_*v*_ ∈ {0, 1}^*m*^ and *σ*(*v*) ∈ [*p*].
2. The root *r*(*T*) is labeled such that *a*_*r*(*T*),*j*_ = 0 if mutation *j* is an SNV and *a*_*r*(*T*),*j*_ = 1 if *j* is an SNP.
3. For each mutation *j*, there is at most one edge (*v, w*) ∈ *E*(*T*) in *T* such that *a*_*v,j*_ = 0 and *a*_*w,j*_ = 1.
4. For each mutation *j*, there are at most *k* edges (*v, w*) ∈ *E*(*T*) such that *a*_*v,j*_ = 1 and *a*_*w,j*_ = 0.
5. For edge (*v, w*) ∈ *E*(*T*) such that *a*_*v,j*_ = 1 and *a*_*w,j*_ = 0 for some *j* ∈ [*m*], we have *σ*(*v*) ≠ *σ*(*w*).
6. For any *ℓ* ∈ [*p*], the set of nodes labeled *σ*(*v*) = *ℓ* form a connected subtree of *T*.

We say that a *n* × *m* binary matrix *A* is a constrained *k*-Dollo phylogeny matrix for copy-number clustering *σ* : [*n*] → [*p*] if and only if there exists a constrained *k*-Dollo phylogeny *T* for *A* and *σ*, i.e. *T* has *n* leaves such that each leaf is labeling by a row *a*_*i*_ of A and *σ*(*i*) for some index *i* ∈ [*n*].

The constrained *k*-Dollo model generalizes the infinite sites model [22] and the *k*-Dollo model [29]. Specifically, when the number *p* of clusters is 1, the constrained *k*-Dollo model is equivalent to the infinite sites model. On the opposite extreme, when the number *p* of clusters is equal to the number *n* of cells, i.e. each cell is in a distinct cluster, the constrained *k*-Dollo model is equivalent to the *k*-Dollo model.

### Constrained *k*–Dollo phylogeny problem for read count data

During a scDNA-seq experiment, we do not observe the mutation matrix *A* directly. Instead, we observe read counts for each mutation in each cell. Specifically, we obtain the variant read count matrix *Q* ∈ ℤ^*n*×*m*^, where *q*_*i,j*_ is the number of reads with the variant allele for mutation *j* in cell *i*, and the total read count matrix *R* ∈ ℤ^*n*×*m*^, where *r*_*i,j*_ is the total number of reads for mutation *j* in cell *i*. Considering that the cells and mutations in each cell are sequenced independently, the likelihood of observing the variant read count matrix *Q* for given total read count matrix *R* and mutation matrix *A* can be written as follows.

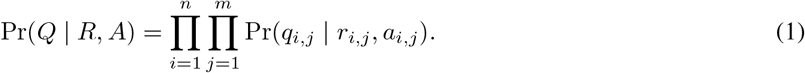

We model the observed variant read counts *q*_*i,j*_ using a beta-binomial, similar to previous work [33, 43, 44]. The “Methods” section provides the details about the read count model.

For given read count matrices *Q* and *R*, copy-number clustering *σ* of the cells and integer *k*, our goal is to construct a constrained *k*-Dollo phylogeny that maximizes the likelihood described in Equation 1. We refer to this as the *Constrained k-Dollo phylogeny problem for read count data* and pose it as follows.

#### Problem 1

(Constrained *k*-Dollo phylogeny problem for read count data (C*k*DP-RC)). Given a variant read count matrix *Q*, total read count matrix *R*, copy-number clustering *σ* and integer *k*, find mutation matrix *A* and phylogeny *T* such that (i) likelihood Pr(*Q* | *R, A*) is maximized and (ii) *T* is a constrained *k*-Dollo phylogeny for *A* and *σ*.

In the “Methods” section, we describe a combinatorial characterization constrained *k*-Dollo phylogenies that we incorporate in an efficient mixed linear integer program (MILP) to solve the C*k*DP-RC problem. Our resulting method, ConDoR, is implemented in Python 3 using `Gurobi` [45] (version 9.0.3) to solve the MILP. ConDoR is available at https://github.com/raphael-group/constrained-Dollo.

### Evaluation on simulated data

We compare ConDoR to SCARLET [33], SPhyR [26], SiFit [30] and SCITE [17] on simulated data. We generated simulated data with *n* ∈ {25, 50, 100} cells, *m* ∈ {25, 50, 100} mutations, *p* ∈ {3, 5}copy-number clusters and maximum number of losses *k* ∈ {1, 2, 3}. We used a growing random network [46] to generate a tree *T* with *m* + *p* edges, and assign mutations, copy number states, and cluster assignments to each tree, as described in the “Methods” section. Next, we assign *n* cells uniformly at random to one of the nodes in the tree. We simulate the sequencing data for each mutation in each cell using a beta-binomial read count model (details in the “Methods” section). We simulate 5 instances for each combination of the varying simulation parameters. The precise input parameters used for each method are described in Additional File 1: Section C.

We compare the mutation matrix *Â* = [*â*_*i,j*_] and tumor phylogeny 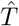 inferred by each method to the ground truth as follows. Following previous studies [26, 33], we evaluate the inferred mutation matrix *Â* against the ground-truth mutation matrix *A* by computing the normalized mutation matrix error *ϵ*(*A, Â*) between *A* and *Â* given by,

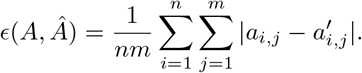

We evaluate the accuracy of the inferred tumor phylogeny 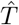 compared to the ground-truth tumor phylogeny *T* by computing the *pairwise ancestral relation accuracy* 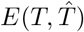 [26, 33]. Specifically, under the assumption that a mutation can be gained only once in the phylogeny, which is used by all methods except SiFit in this study, we compute the accuracy of inferring the correct relationship between all possible pairs of mutations from the inferred tumor phylogeny (details in the “Methods” section). Note that this metric only considers edges of the tumor phylogeny on which mutations are gained and ignores all the edges on which mutations are lost. We exclude SiFit when computing this metric because it a finite-sites model, which allows mutations to occur multiple times in the phylogeny as a consequence of which, pairs of mutations may not have a unique relationship.

ConDoR outperforms all the other methods in terms of both the normalized mutation matrix error (Figure 2a) and the ancestral relationship accuracy (Figure 2b) across all simulation parameters. For instance, on the largest simulated instances with *n* = 100 cells and *m* = 100 mutations, ConDoR achieves the lowest normalized mutation matrix error (median *ϵ*(*A, Â*) = 0.002) and the highest pairwise ancestral relation accuracy (median 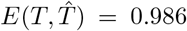) compared to SCARLET (*ϵ*(*A, Â*) = 0.008, 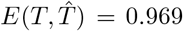), SCITE (*ϵ*(*A, Â*) = 0.01, 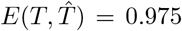), SiFit (*ϵ*(*A, Â*) = 0.05) and SPhyR (*ϵ*(*A, Â*) = 0.02, 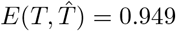). The superior performance of ConDoR comes with running times comparable to existing methods, although ConDoR does have a higher runtime on some of the large simulated instances with *n* = 100 cells and *m* = 100 mutations (Additional File 1: Figure S2).

**Figure 2:**
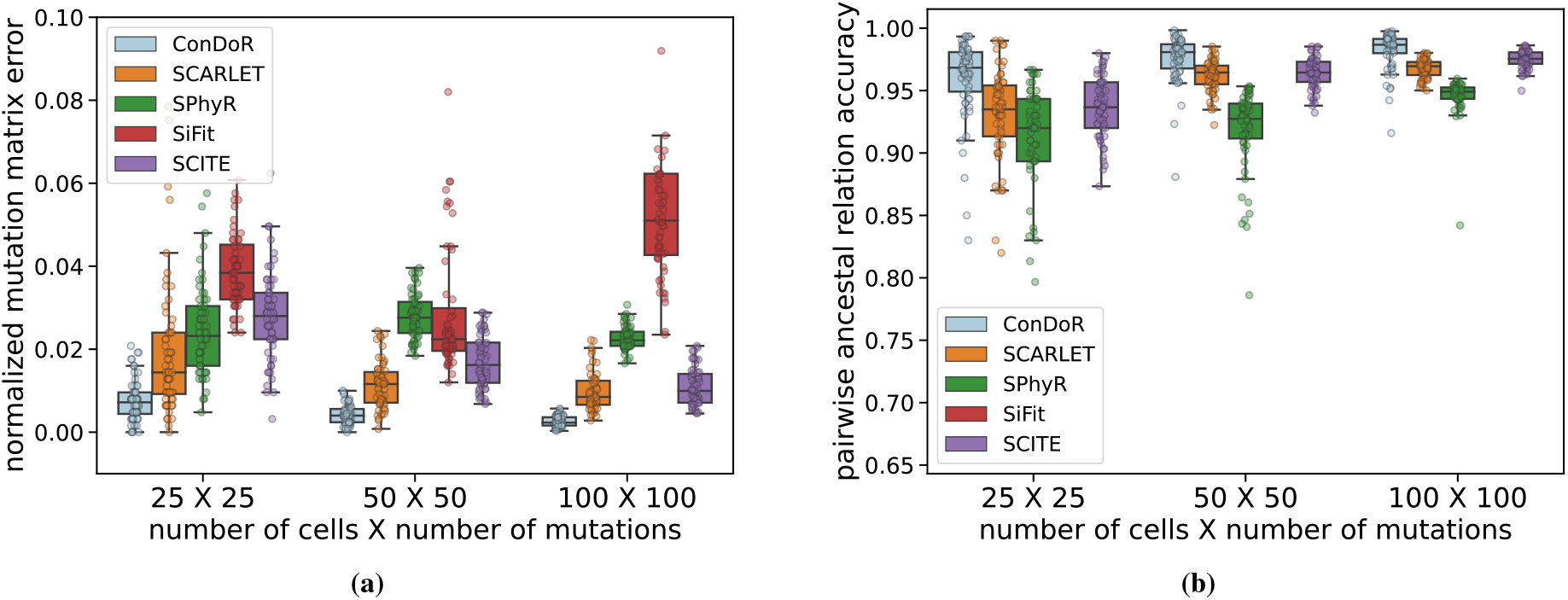
ConDoR outperforms existing methods in recovering the mutation matrix and the tumor phylogeny on simulated data. (a) Normalized mutation matrix error and (b) pairwise ancestral relation accuracy for each method compared to the simulated ground truth. Box plots show the median and the interquartile range (IQR), and the whiskers denote the lowest and highest values within 1.5 times the IQR from the first and third quartiles, respectively.

Interestingly, ConDoR outperforms SCARLET even though SCARLET is given substantially more information about copy number aberrations including both the precise copy-number profile of each cell and the true copy-number tree as input. We believe that this advantage is due to ConDoR solving the underlying optimization problem exactly while SCARLET employs various heuristics that are not guaranteed to yield an optimal solution.

### Multi-region Pancreatic ductal adenocarcinoma data

We used ConDoR to analyze targeted single-cell DNA sequencing (scDNA-seq) data from two regions of a pancreatic ductal adenocarcinoma (PDAC) tumor. Specifically, we sequenced two samples (S1 and S2) from distinct regions of the resected tumor using both conventional bulk whole exome sequencing and Mission Bio Tapestri single-cell sequencing (details in the “Methods” section). The scDNA-seq workflow was conducted using a targeted panel consisting of 596 amplicons (median length is 209 bps, Additional File 1: Figure S3a) interrogating frequently mutated genes in PDAC. We obtained sequencing data from 2153 cells (1167 cells from the first sample and 986 cells from the second sample) with a median coverage of 67× per amplicon per cell.

We identified 7 mutations of interest – including somatic SNVs in *BRCA2, TGFBR2, FGFR1* and germline SNPs in *SPTA1, MGMT*. These mutations were identfied using matched bulk tumor and normal sequencing data and were present in the single-cell data with high confidence (details in “Methods” section). Due to the short length of amplicons and uneven distribution in coverage (Additional File 1: Figure S3b), accurate copy-number calling using this data is challenging. Instead we clustered cells according copy number profiles derived from normalized read counts using *k*-means clustering [47] for number of clusters *p* ∈ {2, …, 8}. We select the best value for *p* using the Silhouette score [48] (see “Methods” section for details). This analysis reveals 3 copy-number clusters (Figure 3a), which we label C0, C1 and C2, that contain 275, 1145 and 733 cells, respectively.

**Figure 3:**
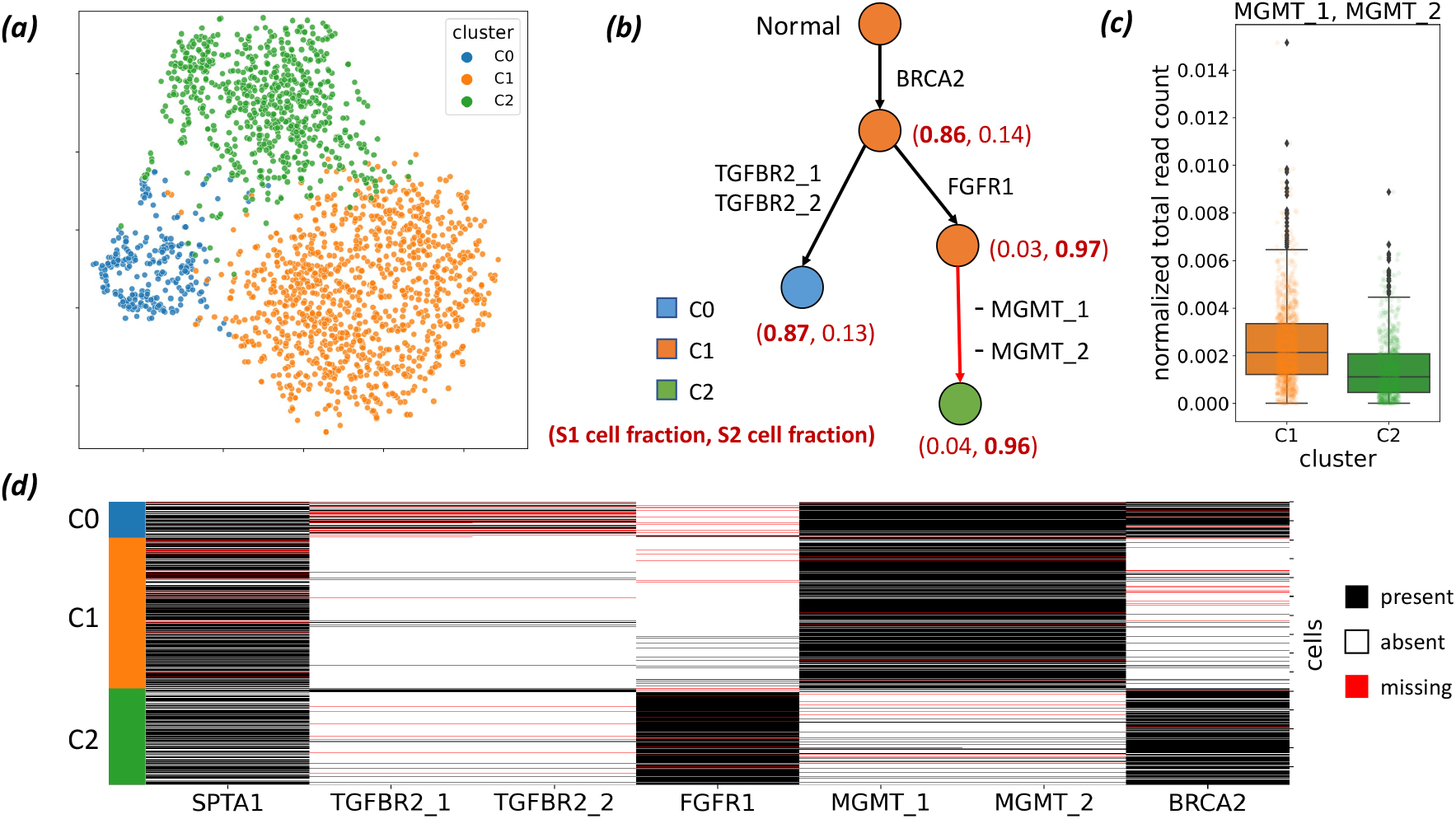
ConDoR provides insights into the evolution and spatial clonal architecture of a pancreatic ductal adenocarcinoma tumor using scDNA-seq data from two different regions of the tumor. (a) *t*-SNE plot showing results of clustering (details in the “Methods” section) of cells into 3 clusters (C0, C1 and C2) according to copy number profiles. (b) Constrained 1-Dollo phylogeny computed by ConDoR with edges labeled by the gain or loss of mutations, and vertices labeled by the copy-number cluster and the fraction of cells from samples S1 and S2 that are attached at that vertex. (c) Reduction in normalized total read count for amplicon AMPL257637 (which contains mutations MGMT_1 and MGMT_2) in cells from cluster C2 compared to cells in cluster C1 (*p* < 5.8 × 10^−33^, a one-sided KS test), supporting the loss of these mutations in the cells belonging to copy-number cluster C2. (d) Observed mutation matrix obtained by discretizing read counts of the 7 mutations, with cells grouped by copy number cluster as indicated in the first column. Box plots show the median and the interquartile range (IQR), and the whiskers denote the lowest and highest values within 1.5 times the IQR from the first and third quartiles, respectively.

ConDoR produces a more plausible phylogeny of the PDAC tumor compared to existing methods and provides insights into the evolution of tumor. While most PDAC cases are driven by canonical gain-of-function *KRAS* mutations [49], ConDoR reveals that the tumor analyzed here is driven by a truncal *BRCA2* stop-gained mutation (p.Y600*), which likely inactivated the BRCA2 protein, a tumor suppressor essential for homologous recombination repair [50]. The ConDoR phylogeny shows branched evolution of the tumor, with the trunk leading to two branches (Figure 3b), the first characterized by two missense *TGFBR2* mutations, TGFBR2_1 (p.A426G) and TGFBR_2 (p.M425I), which likely inactivated cell-intrinsic TGF-*β* signaling, and the second characterized by a missense mutation to *FGFR1* (p.T50K). Although *FGFR1* is involved in MAPK-ERK signaling [49], the particular point mutation’s significance is yet to be characterized. ConDoR infers loss of two germline SNPs in *MGMT*, MGMT_1 and MGMT_2 (both contained in gene MGMT and amplicon AMPL257637), on the edge in the second branch distinguishing cells of cluster C2 from cells of cluster C1. This suggests a loss of heterozygosity (LOH) in cluster C2 cells, which is supported by lower normalized total read count of their amplicon (AMPL257637) in the cells from cluster C2 compared to the cells from cluster C1 (Figure 3c, *p <* 5.8 × 10^−33^ with a one-sided Kolmogorov-Smirnov test). Lastly, the root of the ConDoR phylogeny is labeled by cluster C1, indicating that it contains the normal cells present in the data.

We compared the ConDoR phylogeny to the phylogenies produced by two other methods on this data: COMPASS [35], which infers a comprehensive phylogeny with both SNV and CNA events, and SPhyR [26], which uses the *k*-Dollo model. We could not run SCARLET on this data because it was difficult to obtain reliable integer copy numbers and copy number trees from this targeted sequencing data. While COMPASS takes the read count matrices as input, SPhyR takes an observed mutation matrix (Figure 3d) obtained by discretizing read counts (details in the “Methods section”). COMPASS hypothesizes 8 loss of heterozygosity events (Additional File 1: Figure S4a) covering all genes in the study except *SPTA1* (*BRCA2, TGFBR2, FGFR1, MGMT*). SPhyR (with *k* = 1) produces a phylogeny that contains loss of all the mutations except *FGFR1* (Additional File 1: Figure S4b), which is highly unlikely. This demonstrates that using permissive models, like the ones used in COMPASS and SPhyR, may lead to overfitting of the data resulting in overestimation of mutations with loss. ConDoR’s constrained *k*-Dollo model avoids overfitting by incorporating the copy-number clustering to constrain where loss of mutations can occur in the phylogeny.

The phylogeny constructed by ConDoR also reveals a spatial clonal architecture of the PDAC tumor that agrees with previous histological analysis of the tumor. Specifically, the ConDoR phylogeny shows an enrichment of cells from sample S2 (743 cells from S2 vs. 28 cells from S1) in the second branch of the phylogeny, characterized by the edge with the mutation in *FGFR1* (Figure 3b). Such a spatial separation of the two clonal lineages conforms to histopathological results of this tumor (Figure 5 in [41]) that showed two populations of tumor cells with distinct morphologies that were well demarcated. Spatial structure in the clonal heterogeneity of tumors has also been observed in previous cancer studies and has several clinical implications such as resistance to therapy and recurrences [51–54]. In summary, ConDoR leverages copy-number clustering obtained from targeted scDNA-seq data to build a more plausible tumor phylogeny compared to existing methods and reveals the spatial structure of the intra-tumor heterogeneity.

### Metastatic colorectal cancer data

We also analyzed a published targeted scDNA-seq dataset from a metastatic colorectal cancer patient CRC2 [38]. This dataset consists of 36 SNVs that were identified from a 1000 gene panel in 186 cells; 145 from the primary tumor and 41 from a liver metastasis. The published study build a phylogeny of the 186 cells using SCITE [17] and reported two distinct branches of metastatic cells on this phylogeny. This phylogeny suggests a polyclonal origin of the metastasis, i.e. the metastatic tumor was seeded by two distinct clones that migrated from the primary tumor (Additional File 1: Figure S5a). To evaluate the accuracy of the SCITE tree, the authors identified two *bridge mutations*, in the genes FHIT and ATP7B, that were present in the cells of the second metastatic branch (detected in 10/13 and 13/13 cells, respectively) but absent in the cells of the first metastatic branch (detected in 1/15 and 1/15 cells, respectively).

Two subsequent analyses of this data – using the PhISCS [20] and SCARLET [33] algorithms – yield a simpler explanation for the data; namely that the liver metastasis resulted from monoclonal seeding; i.e. the metastatic tumor resulted from a single migration from the primary tumor. However, neither of these studies adequately explain the absence of the bridge mutations in cells of the second metastatic branch in the SCITE tree. PhISCS removed the bridge mutations from analysis in order to obtain a perfect phylogeny that supports monoclonal seeding. SCARLET, using a loss-supported Dollo model, found evidence for the loss of the FHIT mutation due to a deletion in some cells (Additional File 1: Figure S5b) but concluded that the absence of the ATP7B mutation in all the cells from the second metastatic branch in the SCITE tree was due to simultaneous false negatives in all these cells, a highly unlikely scenario.

ConDoR produces a phylogeny that both supports monoclonal seeding of the metastasis *and* provides a more plausible explanation for the absence of the bridge mutations in some of the metastatic cells compared to previous analyses. The ConDoR phylogeny was produced using the copy-number clustering from [33], which included 4 clusters: 120 diploid cells (D), 33 aneuploid profile of primary tumor cells (P) and two distinct aneuploid profiles of metastatic tumor cells (M1 and M2 with 23 and 10 cells, respectively). The ConDoR tree contains a single branch containing all the metastatic cells, supporting the simpler hypothesis of monoclonal seeding of the liver metastasis, in agreement with the PhISCS and SCARLET analysis (Figure 4a). Moreover, ConDoR infers the loss of both the bridge mutations, FHIT and ATP7B, leading to a phylogeny with a higher likelihood compared to SCARLET (log-likelihood -8324.8 for ConDoR and -8437.4 for SCARLET). This demonstrates that the low resolution of the copy-number aberrations derived from targeted scDNA-seq data used by SCARLET may lead to misleading results, and ConDoR avoids these errors by only using the copy-number clusters.

**Figure 4:**
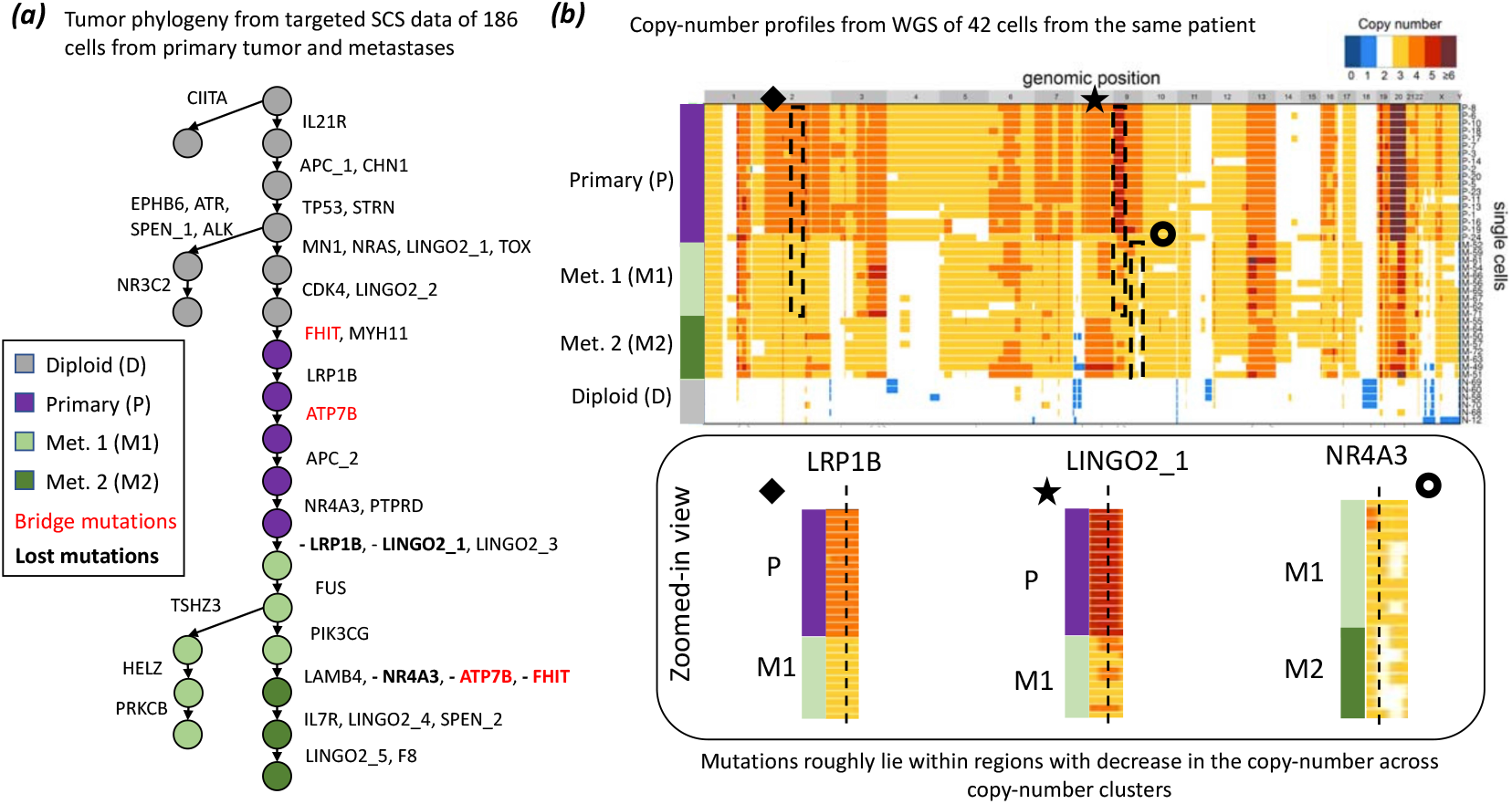
ConDoR infers a phylogeny that is consistent with the copy-number clones in a metastatic colorectal cancer dataset. (a) The ConDoR phylogeny shows loss of bridge mutations FHIT and ATP7B, and suggests monoclonal origin of the liver metastasis. (b) Losses inferred by ConDoR are supported by copy-number profiles from whole genome sequencing data of 42 cells from the same patient in the original study [38] (heatmap showing copy-number profiles adapted from [38]). Mutations LRP1B, LINGO2_1 and NR4A3 lie in regions (black boxes) that decrease in copy-number between the clusters that label the vertices on the edge in the phylogeny where the corresponding mutation ((bold text in (a)) is lost: P → M1 for LRP1B and LINGO2_1, and Ml → M2 for NR4A3.

We also find that the ConDoR tree is consistent with copy-number profiles obtained from whole-genome sequencing of 42 additional cells from the same patient. These cells were were not used in the phylogenetic analyses. In addition to the bridge mutations, ConDoR infers the loss of SNVs in LRP1B, LINGO2 1 and NR4A3. These three SNVs lie within regions with lower copy numbers in the WGS copy-number profiles from the original study (Figure 4b). The copynumber profiles from WGS data also reveal that all metastatic cells share copy-number deletions in chromosomes 2, 3p, 4, 7, 9, 16, and 22 relative to the cells in the primary tumor. These shared copy number profiles further corroborates the ConDoR tree (and the PhISCS and SCARLET trees) in which all metastatic cells are in a single clade. In contrast, SCITE tree from the original study, suggests that these CNAs occurred independently in the two distinct branches of the phylogeny with metastatic cells which is a less likely explanation. In summary, ConDoR integrates SNVs and copy-number clustering to build a tumor phylogeny that contains loss of SNVs that are supported by orthogonal copy-number data and supports a simpler monoclonal origin of the metastasis compared to the original study.

## Conclusions

We introduced a new evolutionary model, the constrained *k*-Dollo model, a model for two-state phylogenetic characters, in which a character can be gained at most once and lost at most *k* times, but where the losses are constrained according to a given clustering of the taxa. This model was inspired by the challenge of inferring a phylogenetic tree from targeted single-cell DNA sequencing data, where SNVs and SNPs are measured with high fidelity, but CNAs are poorly described. Specifically, our model relies on a clustering of cells based on their copy-numbers profiles as input, without requiring identification of precise CNAs in each cell. The constrained *k*-Dollo model generalizes both the infinite sites model and the *k*-Dollo model.

We developed an algorithm, ConDoR (Constrained Dollo Reconstruction), that infers the most parsimonious constrained *k*-Dollo phylogeny using a probabilistic model for the read counts in scDNA-seq data. On simulated data, ConDoR outperforms state-of-the-art tumor phylogeny inference methods. On a multi-region targeted scDNA-seq data of pancreatic ductal adenocarcinoma tumor, ConDoR produced a more plausible phylogeny compared to existing methods, providing insights into the evolution and spatial clonal architecture of the tumor. On targeted scDNA-seq data of metastatic colorectal cancer patient, ConDoR found a phylogeny that supports a simpler monoclonal origin of liver metastasis compared to polyclonal seeding proposed by the original study [38].

There are several limitations and directions for future research. First, ConDoR currently takes the copy-number clustering as input to build a constrained *k*-Dollo phylogeny. A future extension of ConDoR could perform joint inference of the copy-number clustering and the phylogeny, potentially improving the accuracy of both. Second, ConDoR and several existing methods [17, 20, 26, 30, 33] disregard the location of SNVs during phylogeny inference. However, since CNAs alter the copy-number of contiguous segments of the genome, the SNV locations can be used to model the likelihood of simultaneous loss of multiple adjacent SNVs. Lastly, while ConDoR only uses scDNA-seq data as input, the underlying constrained *k*-Dollo model is a general model for evolution of SNVs. We propose that this model can be used for phylogeny inference while integrating information from multiple sequencing technologies, possibly measuring different modalities of the cancer cells [19, 55].

## Methods

### Characterization of constrained *k*-Dollo phylogenies

We derive a characterization of constrained *k*-Dollo phylogeny matrices by building on previous work on characterization of *k*-Dollo phylogeny matrices [26, 27]. Recall that in the *k*-Dollo model, a 0 entry in the mutation matrix *A* indicates that either the mutation did not occur in the cell or that the mutation occurred by then was subsequently lost in the cell. If we could distinguish these two cases, then we could replace the 0 entries resulting losses by additional character states {2, …, *k* + 1} representing the *k* possible losses of a mutation in the *k*-Dollo phylogeny. This idea forms the basis of the following definition of *k-completion of a mutation matrix A*

#### Definition 2

(El-Kebir 2018 [26]). A matrix *B* ∈ {0, …, *k* + 1}^*n*×*m*^ is a *k****-completion*** of a mutation matrix *A* ∈ {0, 1}^*n*×*m*^ provided: (1) *b*_*i,j*_ = 1 if and only if *a*_*i,j*_ = 1; (2) *b*_*i,j*_ ∈ {0, …, *k* + 1} \ {1} if and only if *a*_*i,j*_ = 0; (3) *b*_*i,j*_ ≥ 1 if *j* is an SNP.

The following definition defines a subset of all possible *k*-completion matrices of a mutation matrix *A*.

#### Definition 3

(El-Kebir 2018 [26]). A matrix *B* ∈ {0, …, *k* + 1} ^*n*×*m*^ is a *k****-Dollo completion*** of mutation matrix *A* provided it is a *k*-completion of mutation matrix *A* such that there exists no two columns and three rows in *B* of the following forms:

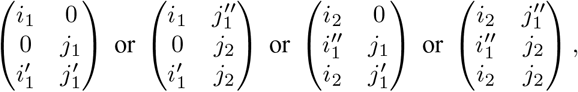

where *i*_1_, 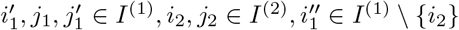 and 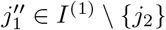, and *I*^(*i*)^ = {*i*, …, *k* + 1}.

According to this definition, the number of 3 × 2 submatrices that are forbidden to exist in *k*-Dollo completion matrices is (*k* + 1)^4^ + 2*k*^2^(*k* + 1)^2^ + *k*^4^. [27] provided an alternate characterization of *k*-Dollo completion matrices, which we describe in Additional File 1: Section B.

*k*-Dollo completion matrices are useful in characterization of *k*-Dollo phylogeny matrices due to the following theorem.

#### Theorem 1

(El-Kebir 2018 [26]). *A* ∈ {0, 1}^*n*×*m*^ is a *k*-Dollo phylogeny matrix if and only if there exists a *k*-Dollo completion *B* ∈ {0, …, *k* + 1}^*n*×*m*^ of *A*.

Constrained *k*-Dollo phylogenies are a subset of *k*-Dollo phylogenies that satisfy some additional constraints. In particular, a constrained *k*-Dollo completion must be *consistent* with copy-number clustering *σ*, according to the following definition.

#### Definition 4

(Consistency). A *k*-Dollo completion *B* ∈ {0, …, *k* + 1} ^*n*×*m*^ of a mutation matrix *A* is *consistent* with a copy-number clustering *σ* with *p* clusters provided the following conditions are true for every mutation *j*.

1. There is at most one cluster *ℓ* such that for two distinct cells *i, i*^′^ ∈ *σ*^−1^(*ℓ*), *b*_*i,j*_ = 0 and *b*_*i*′,*j*_ = 1.
2. If there exists cell *i* such that *σ*(*i*) = *ℓ* and *b*_*i,j*_ = *s* for *s* ∈ {2, …, *k* + 1}, then *b*_*i*′,*j*_ = *s* for all *i*^′^ ∈ *σ*^−1^(*ℓ*). Using this definition, we have the following characterization of constrained *k*-Dollo phylogeny matrices.

#### Theorem 2.

A mutation matrix *A* is a constrained *k*-Dollo phylogeny matrix for copy-number clustering *σ* if and only if there exists a *k*-Dollo completion *B* ∈ {0, …, *k* + 1}^*n*×*m*^ of *A* that is consistent with *σ*.

We provide a proof of Theorem 2 in Additional File 1: Section A and show that given a *k*-Dollo completion *B* of mutation matrix *A* that is consistent with *σ*, we can find a constrained *k*-Dollo phylogeny for *A* and *σ* in *O*(*nmk*) time. In addition, we also show the following result on the complexity of the C*k*DP-RC problem (Problem 1).

#### Theorem 3.

The C*k*DP-RC problem is NP-hard, even for *k* = 0.

A proof of Theorem 3 is provided in Additional File 1: Section A.

### ConDoR algorithm for constrained *k*-Dollo model

We formulate a mixed integer linear program (MILP) to solve Problem 1 exactly. Specifically, for given read count matrices *Q* and *R*, copy-number clustering *σ* and integer *k*, the MILP finds a *k*-Dollo completion *B* that is consistent with *σ* and that maximizes the likelihood Pr(*Q* | *R, A*), where *A* is the mutation matrix corresponding to *B*.

The MILP is based on encoding the combinatorial characterization of constrained *k*-Dollo completion matrices described in “Combinatorial characterization and complexity” subsection. We introduce a binary variables *a*_*i,j*_ for cell *i* and mutation *j* to represent the mutation matrix *A*. Further, we introduce binary variables *c*_*ℓ,j,s*_ for cluster *ℓ*, mutation *j* and state *s* ∈ {2, …, *k* + 1} to represent the presence of loss state *s* of mutation *j* in cluster *ℓ*. These binary variables are used to model the entries of the *k*-completion matrix *B* as follows: *b*_*i,j*_ = 1 if *a*_*i,j*_ = 1; *b*_*i,j*_ = *s* if *c*_*ℓ,j,s*_ = 1 and *σ*(*i*) = *ℓ* for *s* ∈ {2, …, *k* + 1}; and *b*_*i,j*_ = 0 otherwise.

Since *b*_*i,j*_ can only attain one value, we enforce the following constraints.

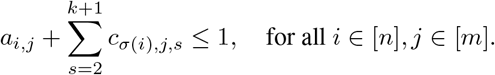

We also define variables *x*_*i,j*_ for cell *i* and mutation *j* which indicates if *b*_*i,j*_ ≥ 1. As such, we enforce

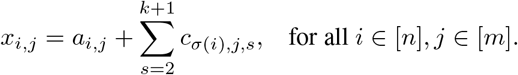

Once we have modeled the *k*-completion matrix *B*, we need to enforce constraints for (i) consistency with the copynumber clustering *σ*, (ii) handling germline mutations and (iii) *B* to be a *k*-Dollo completion matrix. We describe the constraints for (i), (ii) and the objective function of the MILP in the following and refer to Additional File 1: Section B for (iii).

### Handling germline mutations

Here we describe the constraints to handle germline mutations. Note that, if mutation *j* ∈ [*m*] is germline, it must either be present in cell *i* ∈ [*n*], i.e. *a*_*i,j*_ = 1, or it must have been lost, i.e. *c*_*σ*(*i*),*j,s*_ = 1 for some *s* ∈ {2, …, *k* + 1}. As such, if *G* ⊆ [*m*] is the set of germline muations, we enforce the following constraints,

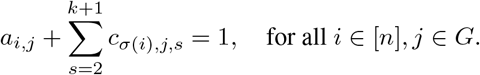

### Consistency constraints

We now describe the constraints to enforce consistency between the *k*-completion matrix *B* and the copy-number clustering *σ*. Note that Condition 2 of Definition 4 is satisfied by the way *B* is modeled and we only need to introduce constraints to satisfy Condition 1 of Definition 4. To that end, we introduce two set of continuous auxiliary variables. First, we introduce 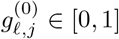 and enforce constraints so that 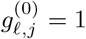 if there exists at least one cell *i* ∈ *σ*^−1^(*ℓ*) such that *b*_*i,j*_ = 0 for cluster *ℓ* and mutation *j*, and 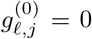 otherwise. Similarly, we introduce 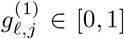 and enforce constraints so that 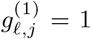 if there exists at least one cell *i* ∈ *σ*^−1^(*ℓ*) such that *b*_*i,j*_ = 1 for cluster *ℓ* and mutation *j*, and 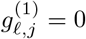 otherwise. We model these variables using the following constraints for all mutations *j* ∈ [*m*] and clusters *ℓ* ∈ [*p*],

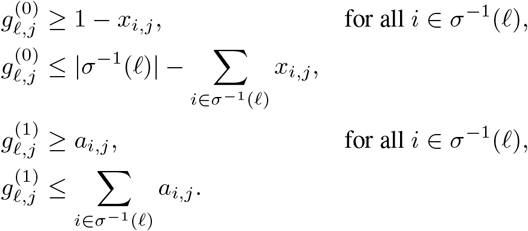

Next, we introduce continuous variables *g*_*ℓ,j*_ ∈ [0, 1] such that *g*_*ℓ,j*_ = 1 if and only if mutation *j* is gained in cluster *ℓ* and *g*_*ℓ,j*_ = 0 otherwise, for cluster *ℓ* and mutation *j*. Specifically, *g*_*ℓ,j*_ = 1 if there exists two distinct cells *i, i*^′^ ∈ *σ*^−1^(*ℓ*) such that *b*_*i,j*_ = 0 and *b*_*i*′,*j*_ = 1. We use 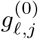 and 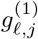 to model *g*_*ℓ,j*_ for all mutations *j* ∈ [*m*] and clusters *ℓ* ∈ [*p*] with the constraints,

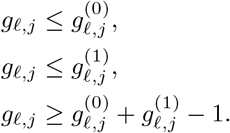

Finally to enforce that each mutation can be gained in at most one cluster, we have the following constraint.

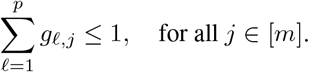

### Objective function

Recall that we want to maximize the likelihood function *P* (*Q|R, A*) (Equation 1), where *A* is the mutation matrix and *B* is its *k*-Dollo completion consistent with copy-number clustering *σ*. After taking log on both sides in Eq. 1, we can linearize the log-likelihood to get the following objective function.

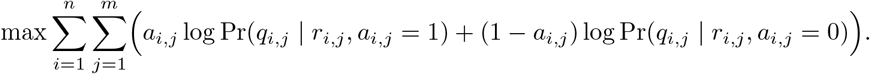

This MILP has *O*(*nm* + *pmk*) binary variables, *O*(*m*^2^*k*^2^ + *pm*) continuous variables, and *O*(*nm*^2^*k*^2^) constraints.

### Simulation details

In this section we provide details about the simulations and the input files generated for each method.

#### Simulation of the phylogeny

We used a growing random network [46] to generate a tree *T* with *m* + *p* edges. Specifically, starting from the root vertex, *T* is built by iteratively adding child nodes while choosing the parent uniformly at random from the nodes in the tree in that iteration. The root node *r*(*T*) represents the normal cell and is assigned to cluster *ℓ* = 0. The edges are then labeled by either the gain of a mutation *j* ∈ [*m*] or change to cluster *ℓ* ∈ [*p*]. For each edge (*v, w*) ∈ *E*(*T*) labeled with a change in cluster, we allow loss of the mutations gained along the path from the root *r*(*T*) to node *v* with probability *λ* = 0.8. We generate a copy-number state for each node in the tree, as described in the “Methods” section.

#### Simulation of copy-number states

SCARLET [33] requires the copy-number profile of each cell as well as the copy-number tree as input. We simulate the copy-number tree as follows.

Each node of the tree is labeled by a copy-number between 0 and 8 for each *j* ∈ [*m*]. We first initialize the root of the tree with a copy-number profile in which the copy-number for each position is picked uniformly at random between 0 and 8. We then label the remaining nodes as we traverse the tree in a breadth-first order. If the edge (*π*(*w*), *w*) does not contain loss of mutation *j*, the copy-number for node *w* is the same as the copy-number of *π*(*w*). On the other hand, if the edge (*π*(*w*), *w*) induces the loss of mutation *j*, the copy-number at *j* for node *π*(*w*) if picked uniformly at random between 1 and 8, while the copy-number of node *w* is picked uniformly at random between 0 and *π*(*w*) − 1. This ensures that, (i) the copy-number profile only changes if there is a loss event on the edge and (ii) each loss of mutation is supported by decrement of copy-number. Let *C* ∈ {0, …, 8} ^*n*×*m*^ be the copy-number matrix such that *c*_*i,j*_ is the copy-number at locus *j* in cell *i*. This copy-number matrix is used during simulation of the variant read counts which we describe in the next subsection.

#### Read count model

The total read count *r*_*i,j*_ for each cell *i* and mutation *j* is modeled as Poisson variable with mean coverage cov = 50.

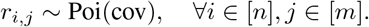

We use beta-binomial model, similar to previous works [33, 43, 44] for the variant read count *q*_*i,j*_ for each cell *i* and mutation *j*. Our model accounts for sequencing errors and allelic imbalance during sequencing as follows. For sequencing error, we set error rate *ϵ* = 0.001 which is similar to the error rates of most recent Illumina sequencing platforms [56]. Specifically, we assume that the false positive rate *α*_*fp*_ and the false negative rate *α*_*fn*_ of observing a read with the variant allele is *ϵ*. When the mutation *j* is not present in cell *i*, i.e. *a*_*i,j*_ = 0, the number of copies of the variant allele is 0. When the mutation *j* is present in cell *i*, i.e. *a*_*i,j*_ = 1, we assume that the number of copies of the variant allele is 1. As such, the value of *a*_*i,j*_ indicates the number of variant allele. Given that the total copies of the locus for mutation *j* in cell *i* is *c*_*i,j*_, the true variant allele frequency, which we denote by *y*_*i,j*_, is given by *y*_*i,j*_ = *a*_*i,j*_*/c*_*i,j*_. Due to sequencing errors *α*_*fp*_ and *ϵ*_*fn*_, the probability *p*_*i,j*_ of producing a read containing the variant allele for mutation *j* in cell *i* is

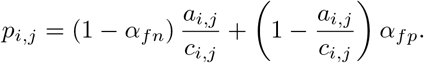

The number of variant reads *q*_*i,j*_ is given by

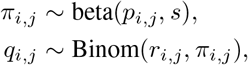

where, we set the dispersion parameter *s* = 15 in our simulations to simulate allelic imbalance. Finally, we spike-in missing entries in the variant read count matrix *Q* and total read count matrix *R* by setting *q*_*i,j*_ = 0 and *r*_*i,j*_ = 0 in ⌊*dnm*⌋ entries where *d* is the rate of missing entries in the data.

While ConDoR and SCARLET take the variant and total read counts as input, several methods (such as SPhyR, SCITE and SiFit) require an observed mutation matrix *A*^′^ as input. In the following section we show how we obtained the observed mutation matrix from the simulated read counts.

#### Obtaining the observed mutation matrix from the read counts

Methods such as SPhyR, SCITE and SiFit take an observed mutation matrix *A*^′^ ∈ {0, 1, − 1} ^*n*×*m*^ as input. This observed mutation matrix *A*^′^ may contain missing entries (represented by − 1) and errors (false positives and false negatives). The aforementioned methods estimate the true binary mutation matrix *A* and build a tumor phylogeny while correcting the errors and imputing the missing entries in the observed mutation matrix *A*^′^. We denote the estimated mutation matrix by *Â*.

We obtain *A*^′^ from the read count matrices *Q* and *R* as follows. We use three filtering parameters to discretize the read count matrices: (i) total read count threshold *r*_*t*_ = 10, (ii) variant read count threshold *q*_*t*_ = 5 and (iii) variant allele frequency threshold *y*_*t*_ = 0.1. We say that mutation *j* is present in cell *i* if and only if the total read count *r*_*i,j*_ is greater than or equal to *r*_*t*_, the variant read count *q*_*i,j*_ is greater than or equal to *q*_*t*_ and the observed variant allele frequency 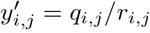 is greater than or equal to *y*_*t*_. Specifically, we set 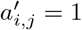 if 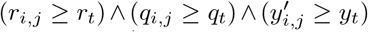. For the remaining entries, we set 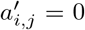 if *r*_*i,j*_ ≥ 0, indicating absence of mutation, and 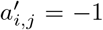, indicating missing entry, otherwise.

#### Pairwise ancestral relationship accuracy

Under the assumption that a mutation can be gained only once in the phylogeny any pair (*j, j*^′^) of mutations can be related in exactly one of the following four ways.

1. Mutation *j* occurs along the path from the root to source node of edge on which mutation *j*^′^ occurs.
2. Mutation *j*^′^ occurs along the path from the root to source node of edge on which mutation *j* occurs.
3. Mutation *j* and *j*^′^ occur on the same edge of the phylogeny.
4. Mutation *j* and *j*^′^ occur on distinct branches of the phylogeny.

We compute the accuracy of inferring the correct relationship between all possible pairs of mutations from the inferred tumor phylogeny.

### Generation and pre-processing of the PDAC data

Here, we provide details regarding the generation and pre-processing of targeted sequencing data of pancreatic ductal adenocarcinoma tumor used in this study.

#### Bulk WES library preparation, sequencing and variant calling

Genomic DNA was extracted from each tissue using the phenol-chloroform extraction protocol [57] or the QIAamp DNA Mini Kits (Qiagen) [58]. WES library preparation and sequencing were performed by the Integrated Genomics Operation at Memorial Sloan Kettering Cancer (MSKCC, NY). Briefly, an Illumina HiSeq 2000, HiSeq 2500, HiSeq 4000 or NovaSeq 6000 platform was used to target sequencing coverages of *>* 250× for WES samples.

The raw FASTQ files were processed with the standard pipeline of the Bioinformatics Core at MSKCC. Sequencing reads were analyzed in silico to assess quality, coverage, and aligned to the human reference genome (hg19) using BWA [59]. After read de-duplication, base quality recalibration and multiple sequence realignment were completed with the PICARD Suite [60] and GATK v.3.1 [61]; somatic single-nucleotide variants and insertion–deletion mutations were detected using Mutect v.1.1.6 [62] and HaplotypeCaller v.2.4 [63]. This pipeline generated a set of mutations for every single sample. Then, all mutations of all samples of the same sequencing cohort were pooled as a single set. Each sample’s BAM file was used to compute “fillout” values (total depth, reference allele read counts, alternative allele read counts) for each mutation in the pooled list. Mutation with alternate read count less than 2 across all samples were removed to trim down false positives. The purpose was to rescue mutations that were detected with high confidence in one sample but with low confidence in another sample of the same patient/tumor. This generated the final output in mutational annotation format (MAF).

#### Single-cell DNA seuqencing (Tapestri) library preparation, sequencing and variant calling

Single nuclei were extracted from snap frozen primary patient samples embedded in optimal cutting temperature (OCT) compound using the protocol developed by Zhang *et al*. [41].

Nuclei were suspended in Mission Bio cell buffer at a maximum concentration of 4000 nuclei/*μ*l, encapsulated in Tapestri microfluidics cartridge, lysed and barcoded. Barcoded samples were then put through targeted PCR amplification with a custom 596-amplicon panel covering important PDAC mutational hotspots in our sample cohort (Table will all the amplicons available at https://github.com/raphael-group/constrained-Dollo).

The 596-amplicon panel was designed based on curation of bulk whole exome/genome sequencing data of PDAC samples collected by the Iacobuzio lab. The goal was to cover as many PDAC-related SNVs within our patient cohort as possible within a 600-amplicon limit, which was deemed economically optimal. The genes/SNVs of interest were determined by querying several resources, such as cBioportal [64, 65] and openCRAVAT [66]. Particular interest was paid to genes in the *TGFβ* pathway as relevant mutations are currently being investigated as clinical biomarkers [67]. In addition to the SNVs, we added amplicons to cover as much exon region as possible for genes that are of particular interest for CNV analyses in PDAC: *KRAS, TP53, SMAD4, CDKN2A, TGFBR1, TGFBR2, ACVR1B, ACVR2A, BMPR1A, BMPR1B, SMAD2, SMAD3, MYC, GATA6, BAP1, MUS81* and *KAT5*.

PCR products were removed from individual droplets, purified with Ampure XP beads and used as templates for PCR to incorporate Illumina i5/i7 indices. PCR products were purified again, quantified with an Agilent Bioanlyzer for quality control, and sequenced on an Illumina NovaSeq. The minimum total read depth was determined by same formula as used in [41].

As described in [41], FASTQ files for single-nuclei DNA libraries were processed through Mission Bio’s Tapestri pipeline with default parameters to arrive at the output H5 file, which mainly consists of two matrices: a single-cell-by-per-amplicon-read-count matrix X_1_, and a single-cell-by-SNV matrix X_2_. Briefly, the pipeline has the following steps.

1. We trim adaptor sequences and align the reads to the hg19 genome (UCSC).
2. We assign reads to individual barcodes, filters for high-quality barcodes. For each of these barcodes, for each amplicon, count the number of forward reads sequenced. This forms matrix X_1_.
3. We generate gVCF for each high-quality barcode’s corresponding BAM file.
4. We jointly genotype gVCF’s of all high-quality barcodes. This forms matrix X_2_.

A more detailed documentation of the pipeline is available at: https://support.missionbio.com/hc/en-us/categories/360002512933-Tapestri-Pipeline. In respect of Mission Bio’s request, the pipeline code is not to be publicized because it contains proprietary information per industry standard. However, the pipeline used in the paper that demonstrated this scDNA-seq library preparation technology [68] is publicly available as a Github repository at https://github.com/AbateLab/DAb-seq. Although we have not formally tested that it performs identically as the Mission Bio pipeline, we believe it is sufficient to replicate our results.

### Variant calling

We detect 40 mutations in the bulk tumor sample with a variant allele frequency (VAF) of at least 0.05. Out of these 40 mutations, 34 mutations were also detected in the matched normal sample indicating that they were germline mutations. From the remaining 6 somatic mutations, we filter out mutations with low prevalence in the scDNA-seq data. Specifically, we only include mutations with variant allele frequency more than 0.1, read depth of more than 20 and variant read depth of more than 10 in at least 5% of the cells. We end up with 4 somatic mutations: `chr3:30715617:C/G` (TGFBR2_1), `chr3:30715619:G/T` (TGFBR2 2), `chr8:38314915:G/T` (FGFR1) and `chr13:32907415:T/A` (BRCA2).

Most phylogeny inference methods only consider somatic SNVs as input, and filter out all germline SNPs. However, germline SNPs that have undergone loss in a subset of cells are informative during phylogeny inference. We identify germline SNPs with putative loss by including SNPs with variant allele frequency less than 0.1, variant read depth more than 10 and total read depth less than 20 in at least 15% of the cells. We find 3 such SNPs: `chr10:131506283:C/T` (MGMT_1), `chr10:131506192:C/T` (MGMT_2) and `chr1:158612236:A/G` (SPTA1). In summary, we consider 3 germline SNPs and 4 somatic SNVs in our analysis.

### Copy-number clustering

In this section we describe the method to cluster the PDAC cells based on the total reads in each cell. Let 𝒜 be the set of amplicons, 𝒢 be the set of genes and 𝒜 (*g*) denote the set of amplicons contained in gene *g* ∈ 𝒢. Let 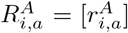 be the *n* × | 𝒜 | *read depth matrix* that contains the number of reads in cell *i* and amplicon *a*. We start by normalizing the amplicon-level read depth matrix by the total reads in each cell. Specifically, we form matrix 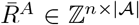 such that,

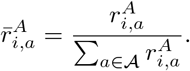

Next, we assume that all loci in the same gene have the same copy-number. As such, we compute the average normalized read depth as follows

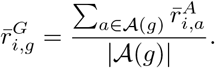

This step helps nullify some noise in the amplicon-level total read count data. We focus on 30 genes with the highest number of amplicons and perform k-means clustering on the resulting matrix 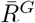.

We use the Silhouette score to determine the number of copy-number clusters. Additional File 1: Figure S6a shows the the Silhouette score for increasing number of clusters in the k-means clustering. We choose the clustering with the highest Silhouette score resulting in *p* = 3 clusters as the copy-number clustering *σ*. Additional File 1: Figure S6b shows a t-SNE [69] where each point is a cell labeled by the cluster index from the copy-number clustering *σ*.

## Acknowledgements

The authors would like to thank the members of the Raphael lab for their helpful discussions in the development of this project. The authors are also grateful to the Single Cell Research Initiative (SCRI), Integrated Genomics Operations (IGO), and the bioinformatics core at MSKCC for their excellent technical support.

## Availability of data and materials

ConDoR is publicly available at https://github.com/raphael-group/constrained-Dollo with the simulation and processed real data analyzed in the paper.

## Ethics approval and consent to participate

Use of samples used in this study was approved by IRB review at Memorial Sloan Kettering Cancer Center.

## Consent for publication

All authors have consented to the publication of this work in Genome Biology.

## Authors’ contributions

P.S. and B.J.R. conceived the project. P.S. characterized the combinatorial structure of the problem, formulated the MILP, and implemented ConDoR. P.S. performed the experimental evaluation. H.Z. performed the sequencing experiments and designed the bioinformatics pipeline for the bulk and the single-cell Tapestri data. All authors interpreted the results. C.A.I-D. and B.J.R. provided supervision. All authors wrote the manuscript. All authors read and approved the manuscript.

## Additional Files

**Additional file 1 — Figures, tables and text describing additional information such as proofs of theorems or additional results**

