## Additional File 1 for "ConDoR: Tumor phylogeny inference with a copy-number constrained mutation loss model"

**C Method parameters** **12**

**D Supplementary results** **13**

### A Proofs

#### A.1 Proof to Theorem 2 in the Main text

Before we start with the proof, we state a few definitions and results that would be useful in the proof.

**Definition 1** ($k$-Dollo state tree, El-Kebir 2018 [1])**.** The $k$-Dollo state tree $S[k]$ is a state tree with nodes $\{0, \ldots, k+1\}$ and edges $\{0, 1\} \cup \{(1, s) | s \in \{2, \ldots, k+1\}\}$.

**Definition 2** (Binary factorization)**.** Let $B$ be a $n \times m$ multi-state character matrix for $m$ characters with $r+1$ states $\{0, 1, \ldots, r\}$ and state tree $S$ with 0 as the root state. The binary factorization $B' = [b'_{i,(j,s)}]$ of $B$ is a $n \times mr$ binary matrix in which $b'_{i,(j,s)} = 1$ for $j \in [m], s \in [r]$ if $b_{i,j} = s'$ where $s \preceq_S s'$, i.e. $s'$ is in the subtree of $S$ rooted at $s$.

The following theorem from [1] relates the $k$-Dollo state tree with $k$-Dollo completion matrices.

**Theorem 1** (El-Kebir 2018 [1])**.** Let $B \in \{0, \ldots, k+1\}^{n \times m}$ be a $k$-Dollo completion matrix. Then there exists a multi-state perfect phylogeny $T_B$ for $B$ such that each mutation $j \in [m]$ has the $k$-Dollo state tree $S[k]$.

The following theorem describes two characterizations of matrices that admits a binary perfect phylogeny.

**Theorem 2** ( [2])**.** Binary matrix $A$ admits a perfect phylogeny if and only if no pair of columns $(j, j')$ that contain the three gametes $(0, 1), (1, 0)$ and $(1, 1)$.

Now, we provide the proof for Theorem 2 in the main text. We restate the theorem here for completeness and provide the proof in the following.

**Theorem 3.** There exists a constrained $k$-Dollo phylogeny $T$ with at most $k$ SNV losses for $A \in \{0, 1\}^{n \times m}$ and copy-number clustering $\sigma : [n] \to [p]$ if and only if there exists a $k$-Dollo completion $B \in \{0, \ldots, k+1\}^{n \times m}$ of $A$ that is consistent with $\sigma$.

*Proof.* ($\Rightarrow$) Let $T$ be a constrained $k$-Dollo phylogeny $T$ for binary matrix $A$ and copy-number clustering $\sigma$ with at most $k$ losses. We construct a $k$-Dollo completion matrix $B$ of $A$ and show that it is consistent with $\sigma$.

We construct $B$ as follows. For SNV $j \in [m]$, let edges of $T$ that induce a loss in SNV $j$ be given by the set $\{e_{j,2}, \ldots, e_{j,q}\}$, where $q \leq k+1$ and the edges are indexed in a breadth first manner. For each leaf

$v \in L(T)$ corresponding to row $i$ in matrix $A$, we set $b_{i,j} = s$ if the edge $e_{j,s}$ is in the unique path from the root $r(T)$ to $v$. For each $(i, j) \in [n] \times [m]$, if $a_{i,j} = 1$, we set $b_{i,j} = 1$. For the rest of the entries we set $b_{i,j} = 0$. It is easy to see that $B$ is a $k$-completion of $A$.

Let us assume that $B$ is not consistent with copy-number clustering $\sigma$. Then for some $j \in [m]$, either condition (1) or condition (2) of Definition 4 in the main text is violated.

**Case 1:** Lets say condition (1) is violated. Then, there must exist two clusters $\ell$ and $\ell'$ such that $b_{i,j} = 0$ and $b_{i',j} = 1$ for distinct cells $i, i'$ of these clusters. Since the set of nodes labeled by the clusters $\ell$ and $\ell'$ are disjoint, the mutation $j$ must be gained at least twice in $T$. However, this contradicts the premise that $T$ is a constrained Dollo phylogeny.

**Case 2:** Lets say condition (2) is violated. Then, there must exist two rows $i$ and $i'$ such that $b_{i,j} = s$ for $s \in \{2, \ldots, k+1\}$, $b_{i',j} \neq s$ and $\sigma(i) = \sigma(i') = \ell$. Let $v$ and $v'$ be the leaves corresponding to the rows $i$ and $i'$ of $B$. Let $r(\ell)$ be the root of the subtree induced by the nodes labeled by $\ell$ in $T$. Consider the path from the root $r(T)$ to $v_i$. The node $r(\ell)$ be lie of this path. As such, we can denote this path by $P = \{r(T), \ldots, r(\ell), \ldots, v_i\}$. Since $b_{i,j} = s$ and $s \in \{2, \ldots, k+1\}$, by construction, there must exist an edge $e \in P$ that induces a loss in SNV $j$. Moreover, since losses can only occur between nodes of $T$ with distinct cluster labeling, edge $e$ most lie in the path $\{r(T), \ldots, r(\ell)\}$. Let $b_{i',j} = t \neq s$. We have three cases, (i) $t = 0$, (ii) $t = 1$ and (iii) $t \in \{2, \ldots, k+1\} \setminus \{s\}$.

**Case (i):** Since $\sigma(v) = \sigma(v') = \ell$ and $T$ is a constrained Dollo phylogeny, $r(\ell)$ must lie in the unique path from $r(T)$ to $v'$. As such, the edge $e$ which induces a loss of SNV $j$ must lie on the path from $r(T)$ to $v'$. However since $b_{i',j} = 0$, by construction, the SNV $j$ is neither lost or gained along the path from $r(T)$ to $v'$, which is a contradiction.

**Case (ii):** Since $b_{i',j} = 1$, by construction, we have $a_{i',j} = 1$. Considering the path from $r(T)$ to $v'$, we have an edge $e$ at which SNV $j$ is lost in the subpath from $r(T)$ to $r(\ell)$. The SNV must be gained in an edge preceding $e$ in the path $P$. Since $a_{i',j} = 1$, the SNV must also be gained in an edge succeeding $e$ in the path $P$. However, this contradicts the premise that $T$ is a constrained Dollo phylogeny in which each SNV can be gained at most once.

**Case (iii):** Since $b_{i',j} = t$ and $t \in \{2, \ldots, k+1\}$, by construction, there must exist an edge $e'$ in the path $P' = \{r(T), \ldots, r(\ell), \ldots, v'\}$ that induces a loss in SNV $j$. Moreover, since losses can only occur between nodes of $T$ with distinct cluster labeling, edge $e'$ must lie in the path $\{r(T), \ldots, r(\ell)\}$. By construction $e$ and $e'$ are distinct since $s \neq t$. Without loss of generality, let $s < t$. As such, the SNV must be gained once along the path $r(T)$ to the source node of $e$ and once along the path from the target node of $e$ to the source node of $e'$. However, this contradicts the premise that $T$ is a constrained Dollo phylogeny in which each

SNV can be gained at most once.

($\Leftarrow$) Let $B$ be a $k$-Dollo completion matrix of $A$ that is consistent with copy-number clustering $\sigma$. For cluster $\ell \in [p]$, let $\sigma^{-1}(\ell)$ be the set of cells in that cluster, i.e.

$$\sigma^{-1}(\ell) = \{i : \sigma(i) = \ell, i \in [n]\}.$$

We show that there exists a constrained $k$-Dollo phylogeny $T$ for $A$ and copy-number clustering $\sigma$. Equivalently, we show that there exists a state tree on the copy-number clusters such that the $k$-Dollo completion matrix $B$ along with the character for the copy-number cluster admits a multi-state perfect phylogeny. Recall that $k$-Dollo completion matrix $B$ admits a $k$-Dollo phylogeny and as such, the binary factorization $B'$ of $B$ with respect to the $k$-Dollo state tree admits a perfect phylogeny.

First, we define a state tree $S$ with the set $\{0, \oplus, 1, 2, \dots, k+1\}$ of states for each character $j \in [m]$ which we will assign to the clusters. The 0 state denotes that mutation $j$ did not occur in the cluster, $\oplus$ denotes that the mutation $j$ was gained in the cluster and $\{1, 2, \dots, k+1\}$ denotes that all cells of the cluster have that state. The state tree $S$ has 0 as the root and has edges $\{(0, \oplus), (\oplus, 1)\} \cup \{(1, s) : s \in \{2, \dots, k+1\}\}$. We will build a character matrix $R \in \{0, \oplus, 1, 2, \dots, k+1\}^{p \times m}$ where each row will represent a cluster as follows.

$$r_{\ell,j} = \begin{cases} s, & \text{if } b_{i,j} = s, \text{for all } i \in \sigma^{-1}(\ell) \text{ and } s \in \{0, 1, 2, \dots, k+1\}, \\ \oplus, & \text{if } b_{i,j} = 0 \text{ and } b_{i',j} = 1, \text{for two cells } i, i' \in \sigma^{-1}(\ell). \end{cases} \tag{12}$$

Since $B$ is consistent with $\sigma$, we can use the above definition to create matrix $R$.

We show that $R$ admits a multi-state perfect phylogeny $T_R$ consistent with the state tree $S$. Consider the binary factorization $R'$ of $R$ with respect to the state tree $S$. We need to show that $R'$ admits a perfect phylogeny. We show this by contradiction. If $R'$ does not admit a binary perfect phylogeny, let $(j, s)$ and $(j', s')$ be two columns of $R'$ that have the three gametes $(0,1), (1,0)$ and $(1,1)$. It is clear that $j \neq j'$ by construction of the binary factorization. Moreover, since each mutation $j$ can be gained in at most one cluster (Condition 1 of Definition 4 in the main text) we have that both $s$ and $s'$ can not be $\oplus$ at the same time. We have the following cases for $s$ and $s'$.

**Case 1:** $s \neq \oplus$ and $s' \neq \oplus$. By construction of $R$, the three gametes $(0,1), (1,0)$ and $(1,1)$, must be present for columns $(j, s)$ and $(j', s')$ in $B'$. But this contradicts the premise that $B$ admits a $k$-Dollo phylogeny.

**Case2:** $s = \oplus$ and $s' \neq \oplus$. Again, since there is at least one cell with $b_{i,j} = 1$ for $i \in \sigma^{-1}(\ell)$ if $r'_{\ell,(j,\oplus)} = 1$, the three gametes $(0,1), (1,0)$ and $(1,1)$, must be present for columns $(j, s)$ and $(j', s')$ in $B'$, which contradicts the premise that $B$ admits a $k$-Dollo phylogeny.

Let $T_R$ define a state tree for a new character $\Sigma$ where each state is a copy-number cluster. We refer to $T_R$ as the copy-number state tree. Let $R''$ be a $n \times p$ binary matrix such that $r''_{i,\ell}$ is given by the binary factorization (Definition 2) of $\Sigma$ associated with node $\sigma(i)$ of the copy-number state tree $T_R$. Note that by construction $R''$ admits a binary perfect phylogeny (it is built by binary factorization of just one multi-state character $\Sigma$). We show that the $n \times (m(k+1)+p)$ augmented matrix $\overline{B} = [B'; R'']$ admits a binary perfect phylogeny by contradiction.

Let us assume that $\overline{B}$ does not admit a binary perfect phylogeny. Then, there must exist two columns such that contain the three gametes $(0,1), (1,0)$ and $(1,1)$. Moreover, one of the columns $\{j^+ : j \in [m]\} \cup \{j_s^- : j \in [m], s \in \{2, \ldots, k+1\}\}$ must be from $B'$ and the other must be one of the columns $[p]$ from $R'$ (since $B'$ and $R'$ both admit a binary perfect phylogeny). There are two possibilities for the column from $B'$.

**Case 1:** Column $(j, s)$ for some $j \in [m], s \in \{2, \ldots, k+1\}$ from $B'$ is in conflict with some column $(\Sigma, \ell)$ for some $\ell \in [p]$ from $R''$. Let $i$ be the cell with gamete $(1,1)$. Since we have gamete $(1,1)$ (i.e. $b'_{i,(j,s)} = 1$ and $r''_{i,(\Sigma,\ell)} = 1$), we have $b_{i,j} = s$ for at least one cell $i \in \sigma^{-1}(\ell)$. Say, gamete $(0,1)$ appears from a cell $i' \in \sigma^{-1}(\ell)$ from the same cluster $\ell$, i.e. $b'_{i',(j,s)} = 0$ and $r'_{i',(\Sigma,\ell)} = 1$. This contradicts the premise that $B$ is consistent with $\sigma$. Alternatively, say gamete $(0,1)$ appears from a cell $i' \in \sigma^{-1}(\ell')$ from a different cluster $\ell'$. This would mean that $\ell'$ is descendant of $\ell$ in $T_R$ (since $r''_{i',(\Sigma,\ell)} = 1$) and $r'(\ell, j) \in \{0, \oplus\}$ (from Eq. 12). However, this contradicts the premise that $T_R$ is a multi-state perfect phylogeny that is consistent with state tree $S$.

**Case 2:** Column $(j, 1)$ for some $j \in [m]$ from $B'$ is in conflict with some column $(\Sigma, \ell)$ for $\ell \in [p]$ from $R''$. Let $i, i'$ and $i''$ be the three cells with gametes $(0,1), (1,1)$ and $(1,0)$. We distinguish two cases.

**Case (i):** $i$ and $i'$ belong to the same cluster $\ell'$ that is descendent of $\ell$ in $T_R$, i.e. $\ell \preceq \ell'$ (since $r''_{i,(\Sigma,\ell')} = r''_{i',(\Sigma,\ell')} = 1$). This would mean that mutation $j$ was gained in cluster $\ell'$, and as such $r'_{\ell,j} = \oplus$. $i''$ must belong to a cluster $\ell''$ that is not descendent of $\ell$ (since $r''_{i'',(\Sigma,\ell)} = 0$). Since mutation can only be gained in at most one cluster, we have $r'_{\ell'',j} = 1$. But $\ell''$ not being descendent of $\ell$ contradicts the premise that $T_R$ is consistent with state tree $S$.

**Case (ii):** $i$ and $i'$ belong to different clusters $\ell$ and $\ell'$. From Eq. 12, we have $r'_{\ell,j} \in \{0, \oplus\}$ and $r'_{\ell,j} \in \{\oplus, 1\}$. Since we have $\ell \neq \ell'$, the possible values for $(r'_{\ell,j}, r'_{\ell',j})$ are $(0, \oplus)$, $(0,1)$ or $\oplus, 1$. Therefore, we have $\ell \prec \ell'$. Again, $i''$ must belong to a cluster $\ell''$ that is not descendent of either $\ell$ or $\ell'$. Since $r''_{i'',(j,1)} = 1$, we have $r'_{\ell'',j} \in \{\oplus, 1\}$. Both options $\oplus$ and $1$ for $r'_{\ell'',j}$ violate the premise that $T_R$ is consistent with state tree $S$ when cluster $\ell''$ that is not descendent of $\ell$.

$\square$

#### A.2 Proof to Theorem 3 in the Main text

We restate the theorem here for completeness and provide a proof in the following.

**Theorem 4.** The C$k$DP-RC problem is NP-hard, even for $k = 0$.

*Proof.* We show this by reduction from the *Flip problem* [3] which is known to be NP-complete and is stated as follows:

**Problem 1** (Flip problem)**.** Given a binary matrix $D$ and an integer $c \in \mathbb{N}$, does there exist a binary matrix $B$ such that (i) $B$ differs from $D$ by at most $c$ entries and (ii) $B$ admits a perfect phylogeny.

Given a instance $(D, c)$ of the Flip problem we construct an instance of the C$k$DP-RC problem with $k = 0$ as follows. We put all the cells $i \in [n]$ in the same copy-number cluster, i.e. $\sigma(i) = 1$ for all $i \in [n]$. The variant read count matrix $Q$ and total read count matrix $R$ as built as follows. For each entry $(i, j) \in [n] \times [m]$ with $d_{i,j} = 0$, we set $q_{i,j} = q_0$ and $r_{i,j} = r_0$, where $q_0$ and $r_0$ are chosen such that $\Pr(q_0 \mid r_0, a_{i,j} = 0) = \mu > 0.5$. For each entry $(i, j) \in [n] \times [m]$ with $d_{i,j} = 1$, we set $q_{i,j} = q_0$ and $r_{i,j} = r_0$, where $q_0$ and $r_0$ are chosen such that $\Pr(q_0 \mid r_0, a_{i,j} = 1) = \mu > 0.5$. Clearly this construction can be completed in polynomial time.

The likelihood of $A$ is given by

$$\Pr(Q \mid R, A) = \prod_{i=1}^{n}\prod_{j=1}^{m}(\Pr(q_1 \mid r_1, a_{i,j}))^{\mathbf{1}(d_{i,j}=1)}(\Pr(q_0 \mid r_0, a_{i,j}))^{\mathbf{1}(d_{i,j}=0)},$$

where $\mathbf{1}(\cdot)$ is the indicator function. Let $c'$ be the number of entries where $D$ and $A$ differ. Then we have

$$\Pr(Q \mid R, A) = \mu^{nm-c'}(1-\mu)^{c'}. \tag{13}$$

We show that the Flip problem instance $(D, c)$ has a solution if and only if the corresponding C$k$DP-RC has a solution with likelihood

$$\Pr(Q \mid R, A) \geq \mu^{nm-c}(1-\mu)^{c}.$$

($\Rightarrow$) Let $B$ be the solution to the Flip problem instance $(D, c)$. From Eq. 13, the likelihood for mutation matrix $A = B$ is given by $\mu^{nm-c}(1-\mu)^{c}$. Moreover, $B$ is a perfect phylogeny and therefore it is also a constrained 0-Dollo phylogeny with the constructed copy-number clustering $\sigma$.

($\Leftarrow$) Let $A$ be the solution to the C$k$DP-RC problem such that

$$\Pr(Q \mid R, A) \geq \mu^{nm-c}(1-\mu)^{c}. \tag{14}$$

Then, suppose $A$ differs from $D$ in $c' > c$ entries. Then from Eq. 13 we have

$$\Pr(Q \mid R, A) = \mu^{nm-c'}(1-\mu)^{c'} < \mu^{nm-c}(1-\mu)^{c},$$

since $\mu > 0.5$. But this contradicts Eq. 14. Hence $A$ must differ from $D$ in at most $c$ entries and thus is a solution to the Flip problem instance $(D, c)$. $\square$

### B MILP formulation

#### B.1 $k$-Dollo completion constraints

Definition 3 in the main text provides $O(k^4)$ distinct $3 \times 2$ submatrices that must be avoided in a $k$-Dollo completion matrix $B$. Incorporating this in the MILP will require $O(n^3m^2k^4)$ constraints which is infeasible for large values of $n, m$ and $k$. We instead employ an alternate characterization of $k$-Dollo completion matrices derived from the work of [4]. In this characterization, for a given matrix $B \in \{0, \ldots, k+1\}^{n \times m}$, we define the $n \times m(k+1)$ *extended binary matrix* $B'$ such that $B$ is a $k$-Dollo completion matrix for some mutation matrix $A$ if and only if $B'$ admits a perfect phylogeny.

**Definition 3.** The extended binary matrix $B' \in 0, 1^{n \times m(k+1)}$ is obtained from the $k$-completion matrix $B$ as follows: for each entry $b_{i,j}$ we have the entry $b'_{i,j^+}$ and $k$ entries $b'_{i,j^-_s}$ (for $s \in \{2, \ldots, k+1\}$) such that (i) if $b_{i,j} = 0$ then $b'_{i,j^+} = 0$ and $b'_{i,j^-_s} = 0$ for $s \in \{2, \ldots, k+1\}$, (ii) if $b_{i,j} = 1$ then $b'_{i,j^+} = 1$ and $b'_{i,j^-_s} = 0$ for $s \in \{2, \ldots, k+1\}$, (iii) if $b_{i,j} = s$ for $s > 1$ then $b'_{i,j^+} = 1$, $b'_{i,j^-_s} = 1$ and $b'_{i,j^-_t} = 0$ for $t \in \{2, \ldots, k+1\} \setminus \{s\}$.

Theorem 3.2 in [5] states the following.

**Lemma 1.** Matrix $B \in \{0, \ldots, k+1\}$ is a $k$-Dollo completion matrix of some mutation matrix $A$ if and only if the corresponding extended binary matrix $B'$ admits a perfect phylogeny.

We now encode the extended binary matrix $B'$ obtained from the $k$-completion matrix $B$ of the mutation matrix $A$ in our MILP formulation. Note that $b'_{i,j^+} = 1$ if and only if $b_{i,j} \geq 1$. As such, the entries $b'_{i,j^+}$ are modeled by the variable $x_{i,j}$. Further, $b'_{i,j^-_s} = 1$ if and only if $b_{i,j} = s$ for $s \in \{2, \ldots, k+1\}$. As such, the entries $b'_{i,j^-_s}$ are modeled by $c_{\sigma(i),j,s}$. We use the *set inclusion and disjointness* (SID) formulation described in [6] to enforce that the extended binary matrix $B'$ is a perfect phylogeny. The SID formulation is based on the following theorem from [2].

**Theorem 5** ( [2]). Let $A = [a_{i,j}]$ be a binary $n \times m$ matrix and let $I(j) := \{i : a_{i,j} = 1\}$ be the set of 1-indices in column $j$. Then $A$ is a perfect phylogeny if and only if for each pair of columns $j$ and $j'$, we have $I(j) \subseteq I(j')$ or $I(j') \subseteq I(j)$ or $I(j) \cap I(j') = \emptyset$.

The above theorem states that a binary matrix admits a perfect phylogeny if and only if for any two columns $j$ and $j'$, their 1-sets (i.e. $I(j)$ and $I(j')$) must be either be related by containment or be disjoint.

Note the extended binary matrix $B'$ has the following $m(k+1)$ columns.

$$\{j^+ : j \in [m]\} \cup \{j_s : j \in [m], s \in \{2, \dots, k+1\}\}$$

. Let the set of loss states $\{2, \dots, k+1\}$ be denoted by $\mathcal{L}_k$. We enforce the conditions described in Theorem 5 on the extended binary matrix $B'$ as follows.

**$k$-Dollo completion constraints** First, we introduce continuous variables that indicate containment between 1-sets of different columns in the extended binary matrix $B'$. We introduce the following set of variables.

1. $y^{(0)}_{j,j',s,s'}$ for $j, j' \in [m]$ and $s, s' \in \mathcal{L}_{||}$, which will be 1 only if 1-set of column $j_s$ contains 1-set of column $j'_{s'}$.
2. $y^{(1)}_{j,j',s'}$ for $j, j' \in [m]$ and $s \in \mathcal{L}_{||}$, which will be 1 only if 1-set of column $j^+$ contains 1-set of column $j'_{s'}$.
3. $y^{(2)}_{j,j',s}$ for $j, j' \in [m]$ and $s \in \mathcal{L}_{||}$, which will be 1 only if 1-set of column $j_s$ contains 1-set of column $j'^+$.
4. $y^{(3)}_{j,j'}$ for $j, j' \in [m]$, which will be 1 only if 1-set of column $j^+$ contains 1-set of column $j'^+$.

We enforce this using the following constraints.

$$y^{(0)}_{j,j',s,s'} \leq 1 - c_{\ell,j',s'} + c_{\ell,j,s}, \qquad \text{for all } \ell \in [p], j, j' \in [m], j \neq j', s, s' \in \mathcal{S}_k, \tag{15}$$

$$y^{(1)}_{j,j',s'} \leq 1 - c_{\ell,j',s'} + \hat{x}_{\ell,j}, \qquad \text{for all } \ell \in [p], j, j' \in [m], j \neq j', s' \in \mathcal{S}_k, \tag{16}$$

$$y^{(2)}_{j,j',s} \leq 1 - (g^{(1)}_{\ell,j'} + \sum_{s'=1}^{k} c_{\ell,j',s'}) + c_{\ell,j,s}, \quad \text{for all } \ell \in [p], j, j' \in [m], j \neq j', s \in \mathcal{S}_k, \tag{17}$$

$$y^{(3)}_{j,j'} \leq 1 - x_{i,j'} + x_{i,j}, \qquad \text{for all } i \in [n], j, j' \in [m], j \neq j'. \tag{18}$$

Next, we introduce variables to indicate if the 1-set of two columns are disjoint.

1. $z^{(0)}_{j,j',s,s'}$ for $j, j' \in [m]$ and $s, s' \in \mathcal{L}_{\|}$, which will be 1 only if 1-sets of column $j_s$ and column $j'_{s'}$ are disjoint.

2. $z^{(1)}_{j,j',s'}$ for $j, j' \in [m]$ and $s \in \mathcal{L}_{\|}$, which will be 1 only if 1-sets of column $j^+$ and column $j'_{s'}$ are disjoint.

3. $z^{(2)}_{j,j'}$ for $j, j' \in [m]$, which will be 1 only if 1-sets of column $j^+$ and column $j'^+$ are disjoint.

We enforce this using the following constraints.

$$
\begin{aligned}
z^{(0)}_{j,j',s,s'} &\leq 2 - c_{\ell,j,s} - c_{\ell,j',s'}, && \text{for all } \ell \in [p], j, j' \in [m], j < j', s, s' \in \mathcal{S}_k, && (19)\\
z^{(1)}_{j,j',s'} &\leq 2 - c_{\ell,j',s'} - (g^{(1)}_{\ell,j} + \sum_{s=1}^{k} c_{\ell,j,s}), && \text{for all } \ell \in [p], j, j' \in [m], j \neq j', s' \in \mathcal{S}_k, && (20)\\
z^{(2)}_{j,j'} &\leq 2 - x_{i,j'} - x_{i,j}, && \text{for all } i \in [n], j, j' \in [m], j < j'. && (21)
\end{aligned}
$$

Following Theorem 5, we require the following conditions to attain a perfect phylogeny.

$$
\begin{aligned}
y^{(0)}_{j,j',s,s'} + y^{(0)}_{j',j,s',s} + z^{(0)}_{j,j',s,s'} &\geq 1, && \text{for all } j, j' \in [m], j < j', s, s' \in \mathcal{S}_k, && (22)\\
y^{(1)}_{j,j',s'} + y^{(2)}_{j',j,s'} + z^{(1)}_{j,j',s'} &\geq 1, && \text{for all } j, j' \in [m], j \neq j', s, s' \in \mathcal{S}_k, && (23)\\
y^{(3)}_{j,j'} + y^{(3)}_{j',j} + z^{(2)}_{j,j'} &\geq 1, && \text{for all } j, j' \in [m], j < j'. && (24)
\end{aligned}
$$

This formulation contains $O(nm^2k^2)$ constraints with $O(m^2k^2)$ continuous variables.

##### B.2 Complete MILP

For completeness sake, we describe the complete MILP here.

$$
\begin{aligned}
\max\ & \sum_{i=1}^{n}\sum_{j=1}^{m}\Big(a_{i,j}\log\Pr(q_{i,j}\mid r_{i,j},a_{i,j}=1)+ \\
& \qquad (1-a_{i,j})\log\Pr(q_{i,j}\mid r_{i,j},a_{i,j}=0)\Big) \\
\text{s.t. } & a_{i,j}+\sum_{s=2}^{k+1}c_{\sigma(i),j,s}\le 1, && \forall i\in[n],j\in[m], \\
& x_{i,j}=a_{i,j}+\sum_{s=2}^{k+1}c_{\sigma(i),j,s} && \forall i\in[n],j\in[m], \\
& g^{(0)}_{\ell,j}\ge 1-x_{i,j}, && \forall i\in\sigma^{-1}(\ell),\ell\in[p],j\in[m], \\
& g^{(0)}_{\ell,j}\le|\sigma^{-1}(\ell)|-\sum_{i\in\sigma^{-1}(\ell)}x_{i,j}, && \forall \ell\in[p],j\in[m], \\
& g^{(1)}_{\ell,j}\ge a_{i,j}, && \forall i\in\sigma^{-1}(\ell),\ell\in[p],j\in[m], \\
& g^{(1)}_{\ell,j}\le\sum_{i\in\sigma^{-1}(\ell)}a_{i,j}, && \forall \ell\in[p],j\in[m], \\
& g_{\ell,j}\le g^{(0)}_{\ell,j},\ g_{\ell,j}\le g^{(1)}_{\ell,j},\ g_{\ell,j}\ge g^{(0)}_{\ell,j}+g^{(1)}_{\ell,j}-1, && \forall \ell\in[p],j\in[m], \\
& \sum_{\ell=1}^{p}g_{\ell,j}\le 1, && \forall j\in[m], \\
& y^{(0)}_{j,j',s,s'}\le 1-c_{\ell,j',s'}+c_{\ell,j,s}, && \forall \ell\in[p],j,j'\in[m],j\ne j',s,s'\in\mathcal{S}_k, \\
& y^{(1)}_{j,j',s'}\le 1-c_{\ell,j',s'}+\hat{x}_{\ell,j}, && \forall \ell\in[p],j,j'\in[m],j\ne j',s'\in\mathcal{S}_k, \\
& y^{(2)}_{j,j',s}\le 1-(g^{(1)}_{\ell,j'}+\sum_{s'=2}^{k+1}c_{\ell,j',s'})+c_{\ell,j,s}, && \forall \ell\in[p],j,j'\in[m],j\ne j',s\in\mathcal{S}_k, \\
& y^{(3)}_{j,j'}\le 1-x_{i,j'}+x_{i,j}, && \forall i\in[n],j,j'\in[m],j\ne j', \\
& z^{(0)}_{j,j',s,s'}\le 2-c_{\ell,j,s}-c_{\ell,j',s'}, && \forall \ell\in[p],j,j'\in[m],j< j',s,s'\in\mathcal{S}_k, \\
& z^{(1)}_{j,j',s'}\le 2-c_{\ell,j',s'}-(g^{(1)}_{\ell,j}+\sum_{s=2}^{k+1}c_{\ell,j,s}), && \forall \ell\in[p],j,j'\in[m],j\ne j',s'\in\mathcal{S}_k, \\
& z^{(2)}_{j,j',s,s'}\le 2-x_{i,j'}-x_{i,j}, && \forall i\in[n],j,j'\in[m],j< j', \\
& y^{(0)}_{j,j',s,s'}+y^{(0)}_{j',j,s,s'}+z^{(0)}_{j,j',s,s'}\ge 1, && \forall j,j'\in[m],j< j',s,s'\in\mathcal{S}_k, \\
& y^{(1)}_{j,j',s'}+y^{(2)}_{j',j,s'}+z^{(1)}_{j,j',s'}\ge 1, && \forall j,j'\in[m],j\ne j',s,s'\in\mathcal{S}_k, \\
& y^{(3)}_{j,j'}+y^{(3)}_{j',j}+z^{(2)}_{j,j'}\ge 1, && \forall j,j'\in[m],j< j',
\end{aligned}
$$

$$
\begin{aligned}
&a_{i,j} \in \{0,1\}, && \forall i \in [n], j \in [m], \\
&c_{\ell,j,s} \in \{0,1\}, && \forall \ell \in [p], j \in [m], s \in \mathcal{S}_k, \\
&g_{\ell,j}, g_{\ell,j}^{(0)}, g_{\ell,j}^{(1)}, \hat{x}_{\ell,j} \in [0,1], && \forall \ell \in [p], j \in [m], \\
&y_{j,j',s,s'}^{(0)}, y_{j,j',s,s'}^{(1)}, y_{j,j',s,s'}^{(2)}, y_{j,j',s,s'}^{(3)} \in [0,1], && \forall j, j' \in [m], j \neq j', s' \in \mathcal{S}_k, \\
&z_{j,j',s,s'}^{(0)}, z_{j,j',s,s'}^{(1)}, z_{j,j',s,s'}^{(2)} \in [0,1], && \forall j, j' \in [m], j < j', s' \in \mathcal{S}_k.
\end{aligned}
$$

### C Method parameters

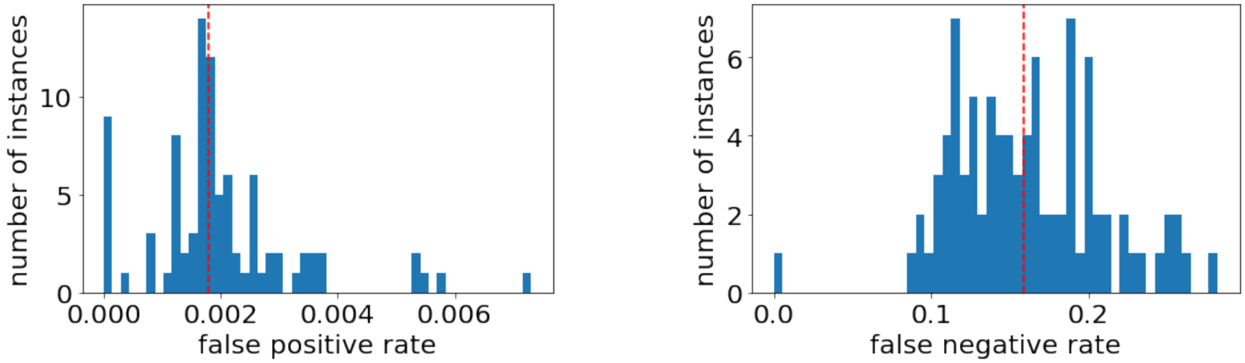

(a) (b)

**Figure S1:** Histogram of the (a) false positive rate $\alpha$ and the (b) false negative rate $\beta$ in the observed mutation matrix $A'$ of simulated instances. Red lines show the median false positive rate (0.0018) and median false negative rate (0.158).

Here we provide the precise commands used to run competing methods for benchmarking on the simulated instances. Methods such as SPhyR, SCITE and SiFit take error rates, i.e. false positive rate and false negative rate, as input during inference. For simulated data, we empirically estimate the false positive rate and false negative rates (Figure S1) in the observed mutation matrix $A'$ obtained in the simulations. We provide the aforementioned methods with the median false positive rate $\alpha = 0.0018$ and median false negative rate $\beta = 0.158$ as input. Description of keywords used in the command-line arguments are provided in Table S1.

**ConDoR** We run ConDoR with both false positive and false negative sequencing error rates set to $0.001$.

```
python condor.py -i ${input_clustering}
-a ${params_fp} -b ${params_fn} -k ${k}
-r ${input_total_readcounts} -v ${input_variant_readcounts}
-o ${output_prefix}
```

**SCARLET** We run SCARLET with the copy-number tree as well as the copy-number profiles from the simulations as input.

```
python2 scarlet.py ${input_combined_readcount_file} ${copy_number_tree_file}
${output_prefix}
```

| | command-line keyword | description |
|---|---|---|
| $n$ | ${num_cells}$ | number of cells |
| $m$ | ${num_mutations}$ | number of mutations |
| $A'$ | ${input_character_matrix} | observed mutation matrix |
| $Q$ | ${input_variant_readcounts} | variant allele read count matrix |
| $R$ | ${input_total_readcounts} | total read count matrix |
| $\sigma$ | ${input_clustering}$ | copy-number clustering |
| $(Q, R)$ | ${input_combined_readcount_file}$ | single file containing both $Q$ and $R$ |
| $k$ | ${k} | maximum losses allowed for any mutation |

**Table S1:** Description of the keywords used in the command-line arguments.

**SPhyR** We run SPhyR with the flag -lT (which sets the number of distinct rows in the inferred mutation matrix) set to $m + p$ since that is the maximum number of events that can occur in the phylogeny.

```
./kDPFC ${input_character_matrix} ${output_filename}
-a 0.0018 -b 0.158 -k ${k}
-lT ${params_lT} -lC ${params_lC}
```

**SCITE** We run SCITE for 1000000 iterations. Since we do not have any homozygous mutations in the simulations, we set -cc flag to 0.

```
scite -i ${input_character_matrix} -n ${num_mutations}$ -m ${num_cells}$
-o ${output_prefix}$ -a -cc 0 -l 1000000 -r 1
-fd 0.0018 -fn 0.158 0 -max_treelist_size 1
```

**SiFit** We run SiFit for 1000000 iterations.

```
java -jar sifit -ipMat ${input_character_matrix}
-m ${num_cells} -n ${num_mutations}$
-df 0 -l 1000000 -r 1 -fp 0.0018 -fn 0.0158
```

### D Supplementary results

We have the following supplementary figures and tables.

- Figure S2 shows the runtime of ConDoR, SCARLET, SPhyR, SiFit and SCITE on the simulation instances ("Evaluation on simulated data" section in the Main text).

- Figure S3 shows the distribution of length and coverage of the 596 amplicons used in the PDAC study ("Multi-region Pancreatic ductal adenocarcinoma data" section in the Main text).
- Figure S4 shows the phylogeny constructed using COMPASS and SPhyR on the PDAC dataset ("Multi-region Pancreatic ductal adenocarcinoma data" section in the Main text).
- Figure S5 shows the phylogeny constructed using SCITE and SCARLET on metastatic CRC data ("Metastatic colorectal cancer data" section in the Main text).
- Figure S6 shows the Silhouette score for increasing number of clusters in the k-means clustering of cells in the PDAC data and the TSNE plot showing the clustering with the best Silhouette score ("Multi-region Pancreatic ductal adenocarcinoma data" section in the Main text).

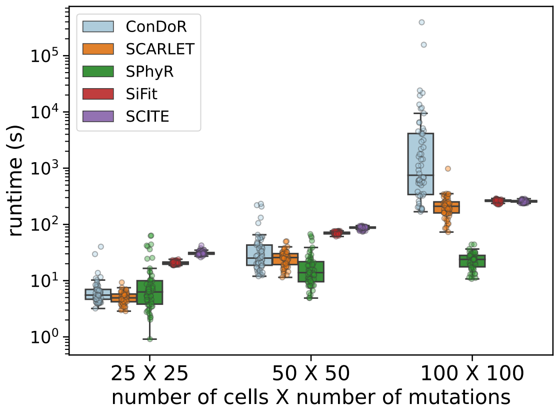

**Figure S2:** Computational time taken by all the methods on simulated instances ("Evaluation on simulated data" section in the Main text).

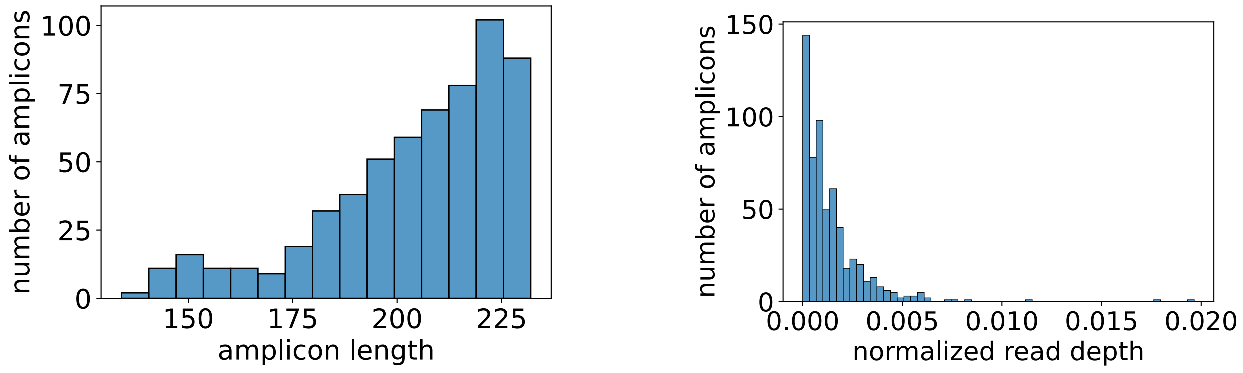

**(a)** **(b)**

**Figure S3:** Histogram of (a) length and (b) normalized read depth for the 596 amplicons used for targeted single-cell sequencing in the PDAC study ("Multi-region Pancreatic ductal adenocarcinoma data" section in the Main text).

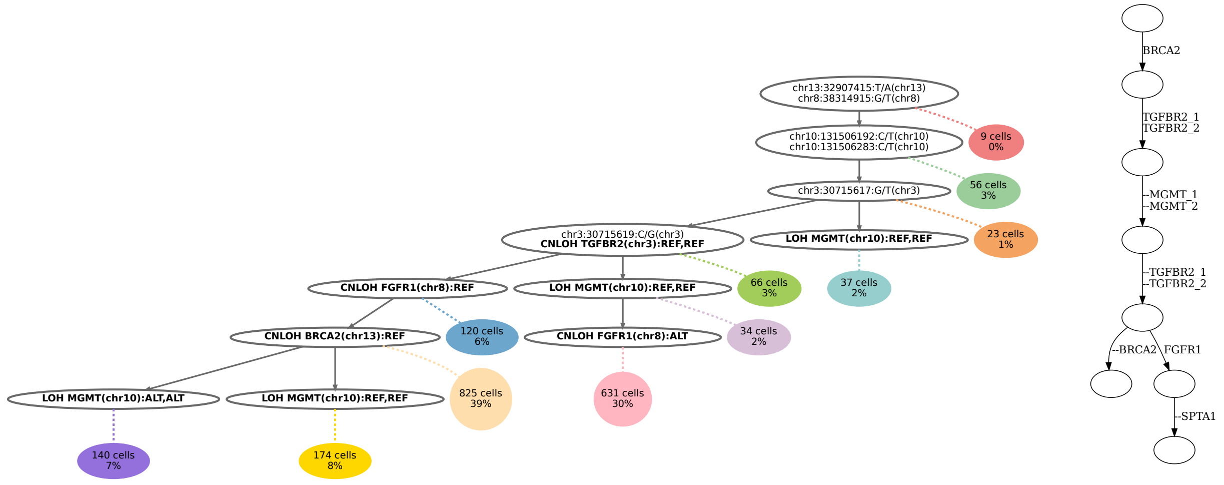

**(a)** **(b)**

**Figure S4:** Phylogeny inferred by (a) COMPASS [7] and (b) SPhyR [1] on the PDAC data. COMPASS infers 8 independent LOH events across 4 genes (MGMT, BRCA2, FGFR1, TGFBR2), while SPhyR infers the loss of 6 out of the 7 mutations ("Multi-region Pancreatic ductal adenocarcinoma data" section in the Main text).

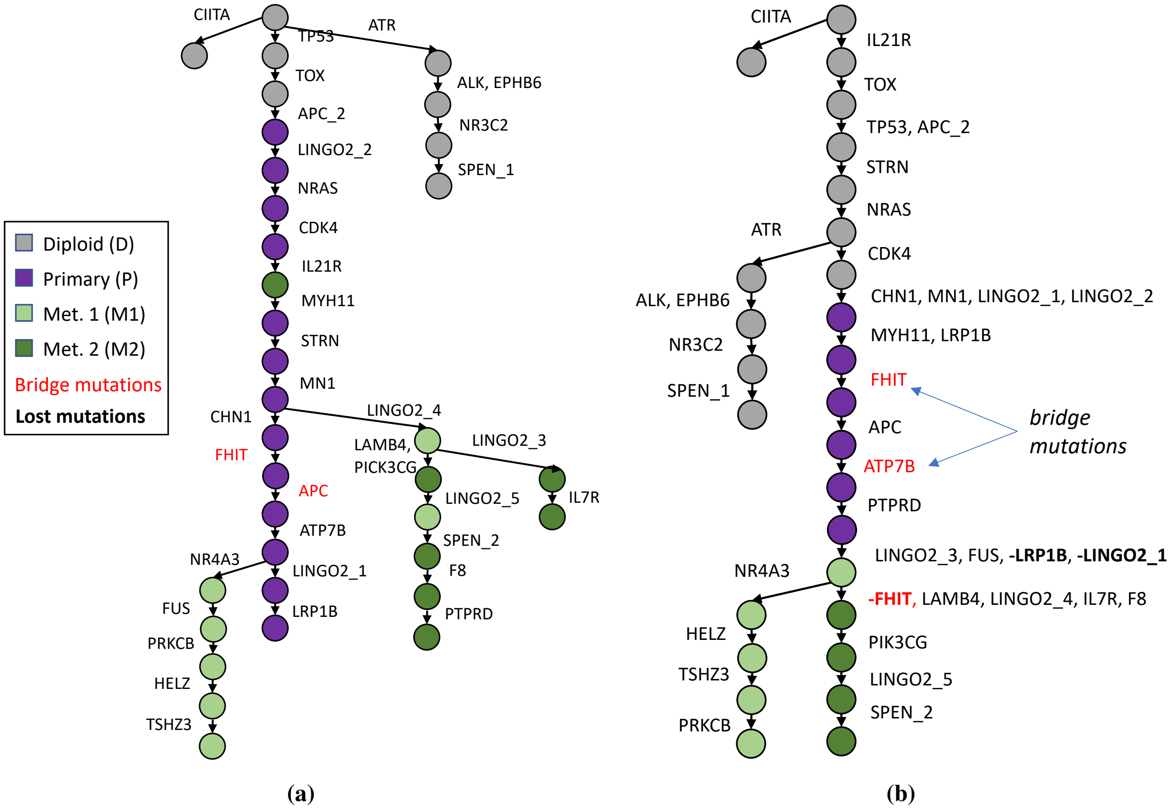

**Figure S5:** Phylogeny inferred by (a) SCITE [8] and (b) SCARLET [9] on the metastatic colorectal cancer data ("Metastatic colorectal cancer data" section in the Main text).

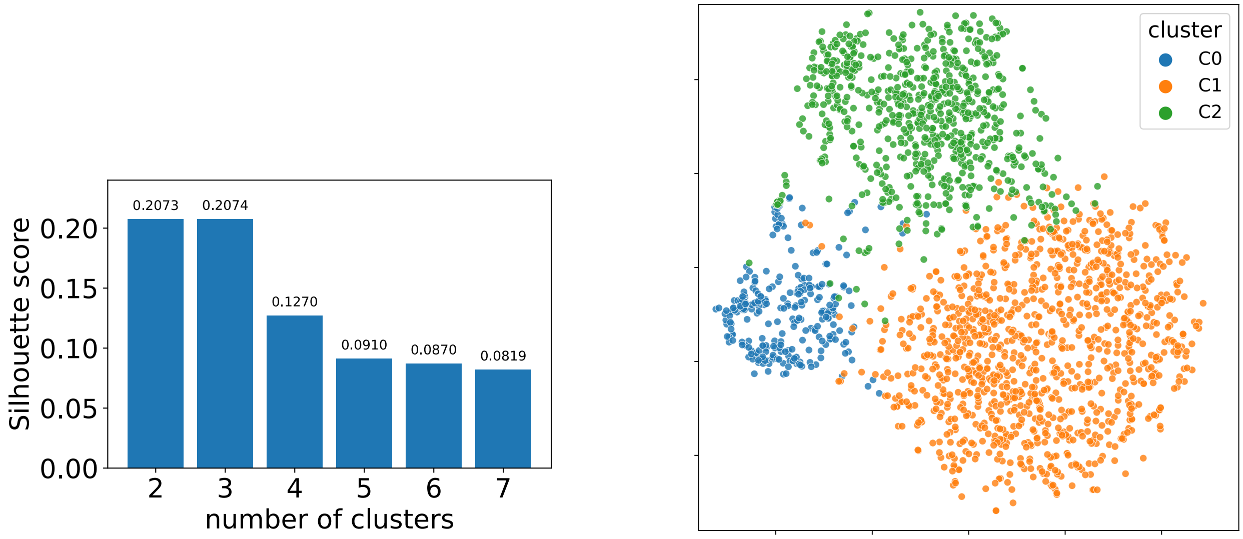

**(a)** **(b)**

**Figure S6:** (a) Silhouette score for increasing number of clusters in k-means clustering of cells using the gene-level average normalized read depth $\bar{R}^G$ for the PDAC dataset. (b) tSNE plot showing the k-means clustering (that has 3 clusters) with the highest Silhouette score (0.96). Related to "Multi-region Pancreatic ductal adenocarcinoma data" section in the Main text.

[7] Etienne Sollier, Jack Kuipers, Niko Beerenwinkel, Koichi Takahashi, and Katharina Jahn. Joint copy number and mutation phylogeny reconstruction from single-cell amplicon sequencing data. *bioRxiv*, 2022.

[8] Katharina Jahn, Jack Kuipers, and Niko Beerenwinkel. Tree inference for single-cell data. *Genome biology*, 17(1):1–17, 2016.

[9] Gryte Satas, Simone Zaccaria, Geoffrey Mon, and Benjamin J Raphael. Scarlet: Single-cell tumor phylogeny inference with copy-number constrained mutation losses. *Cell systems*, 10(4):323–332, 2020.
